# Impact of Concatenating fMRI Data on Reliability for Functional Connectomics

**DOI:** 10.1101/2020.05.06.081679

**Authors:** Jae Wook Cho, Annachiara Korchmaros, Joshua T Vogelstein, Michael Milham, Ting Xu

## Abstract

Compelling evidence suggests the need for more data per individual to reliably map the functional organization of the human connectome. As the notion that ‘more data is better’ emerges as a golden rule for functional connectomics, researchers find themselves grappling with the challenges of how to obtain the desired amounts of data per participant in a practical manner, particularly for retrospective data aggregation. Increasingly, the aggregation of data across all fMRI scans available for an individual is being viewed as a solution, regardless of scan condition (e.g., rest, task, movie). A number of open questions exist regarding the aggregation process and the impact of different decisions on the reliability of resultant aggregate data. We leveraged the availability of highly sampled test-retest datasets to systematically examine the impact of data aggregation strategies on the reliability of cortical functional connectomics. Specifically, we compared functional connectivity estimates derived after concatenating from: 1) multiple scans under the same state, 2) multiple scans under different states (i.e. hybrid or general functional connectivity), and 3) subsets of one long scan. We also varied connectivity processing (i.e. global signal regression, ICA-FIX, and task regression) and estimation procedures. When the total number of time points is equal, and the scan state held constant, concatenating multiple shorter scans had a clear advantage over a single long scan. However, this was not necessarily true when concatenating across different fMRI states (i.e. task conditions), where the reliability from the aggregate data varied across states. Concatenating fewer numbers of states that are more reliable tends to yield higher reliability. Our findings provide an overview of multiple dependencies of data concatenation that should be considered to optimize reliability in analysis of functional connectivity data.

## INTRODUCTION

Functional connectomics is a mainstream modality for mapping brain organization (Greicius et al., 2003; Power et al., 2014; Smith et al., 2013; Sporns, 2013; Yeo et al., 2011). Initially focused on the demonstration of findings at the population-level, the past decade has witnessed a burgeoning literature focused on the delineation of differences among individuals and clinical groups (Braga and Buckner, 2017; Fornito and Bullmore, 2015; Laumann et al., 2015). Recognizing concerns about reproducibility stemming from a reliance on small samples, researchers have raised the bar for sample size, leveraging the growing availability of public datasets to achieve this goal, for example, Human Connectome Project, UK Biobank, 1000 Functional Connectomes, Consortium for Reliability and Reproducibility, etc. (Biobank, 2014; Biswal et al., 2010; Van Essen et al., 2013; X. N. Zuo et al., 2014). Unfortunately, the sample size is not the sole consideration for efforts to achieve reproducible findings in studies of individual and group-differences. The reliability of the measures obtained directly impacts the sample sizes required and the likelihood of replication (Matheson, 2019; Zuo et al., 2019). Recognizing this reality, researchers are increasingly emphasizing the need to obtain larger amounts of data per individual, as reliability appears to be directly related (Birn et al., 2013; Gordon et al., 2015; Noble et al., 2017; Xu et al., 2016).

While the need for more data per individual appears to be straightforward, how to achieve the desired amount of data is not. It is not always feasible to obtain more data due to practical limitations of a crowded scan protocol, or tolerability of the scanning environment (Krause et al., 2019; Meissner et al., 2020; Menon et al., 1997). Additionally, this solution does little to address questions about how to improve the amount of data available for analysis in existing data acquisitions. In response, a growing theme in the literature suggests that fMRI data from different fMRI scan states (e.g., rest, task, movie) can be combined to achieve the desired quantities (Elliott et al., 2019; O’Connor et al., 2017; Schmälzle et al., 2017). At present, there is limited guidance about the relative trade-offs of using concatenated data, or how to best do so.

In this study, we examine the impact of data concatenation strategies on the reliability of functional connectomics and differences that can arise from the specific concatenation method employed. First, we examine the impact of concatenating data from multiple short scans versus a single long scan. This is an important issue in the literature, as studies focused on hyperkinetic populations, such as the Adolescent Brain Cognitive Development (ABCD) study by National Institutes of Health (NIH), have advocated for using multiple shorter scans as a means of improving tolerability and compliance (i.e., low head motion) (Casey et al., 2018). Second, we examine the impact of concatenating data from different states versus the same state. A recent study suggested that the calculation of functional connectivity from multiple states yields data that has superior reliability to that from the same state (Elliott et al., 2019). In this work, they compared a hybrid dataset, generated by concatenating data from different scans versus resting-state data. One question that arises is if the observed advantage was truly the product of combining different states, versus the concatenation of small segments of data from different scans - which was shown to be advantageous (relative to fewer large segments) in the single-subject MyConnectome dataset (Laumann et al., 2015). Third, we address the question of whether there is any advantage to first calculating connectivity matrices for each scan and then combining, or to concatenating time-series data and then calculating connectivity. To date, there is little guidance in the literature regarding this issue. Finally, we examine the impact of the number of test-retest units versus the amount of data per unit on test-retest reliability as well as how concatenation of smaller segments of data contributes to different combinations of numbers of test-retest units and length of each subset (Noble et al., 2017).

It is important to clarify that the goal of the present work is not to question the value of combining all available data in a given study to optimize reliability for a given collection, as that is obvious from the existing literature (Elliott et al., 2019; Noble et al., 2019, 2017; Xu et al., 2016). Instead, our goals are twofold. First, to address the issue of data concatenation and how to understand the trade-offs of different decisions when designing a new experiment for the acquisition of a fixed amount of data. Second, to clarify potential confusions in the literature regarding determinants of reliability - most notably, the reported advantages for hybrid (i.e., mixed state) functional connectivity measures, and the relative trade-offs related to the number of sessions versus scan duration (Elliott et al., 2019; Noble et al., 2017).

## METHODS

All analyses in the present work were carried out using publicly available datasets selected based on the inclusion of extensive test-retests fMRI.

### Human Connectome Project (HCP)

We made use of the HCP S1200 release dataset (Glasser et al., 2013; Marcus et al., 2013). The unrelated healthy young adult participants who completed all fMRI scans (i.e. rest and task) were considered in our study. We selected the low head motion participants (mean FD < 0.25 mm, max FD < 2.75 mm). The final sample used in this study included 217 (104 male, age=28.5 +/- 3.7 years) unrelated participants. Extensive descriptions of the data and the minimal preprocessing applied can be found in prior publications (e.g., (Glasser et al., 2013; Marcus et al., 2013). In brief, for each participant, the dataset includes 4 resting fMRI (rfMRI) scans obtained across two days - namely REST1_LR, REST1_RL, REST2_LR, and REST2_RL in HCP files. Here the LR and RL represent left-to-right and right-to-left phase coding directions that were used in the gradient-echo EPI sequence (TR=0.72s, TE=33.1ms, flip angle=52°, FOV=208×180mm, resolution=2×2×2mm); each rfMRI scan contains 1200 volumes (~15 mins). To control for the effects of head motion, we further selected a subsample from the HCP dataset as a low-motion replication dataset. Specifically, we selected participants with ‘clean’ rfMRI data after the HCP minimal preprocessing on the Power-‘voxplot’ (Power et al., 2017). This subsample consists of 22 subjects from 217 subjects selected by visual inspection (subject ID in Supplementary Table 1).

### HCP test-retest dataset

We also used the HCP test-retest dataset, which contains 7 tasks and 2 resting-state sessions, repeated twice per participant. Six tasks were used in our analyses including working memory (WM, 405 volumes), gambling (GAM, 253 volumes), motor tasks (MOT, 284 volumes), which were collected during the first day, and language (LAN, 316 volumes), social (SOC, 274 volumes) and relational (REL, 232 volumes) tasks collected on the second day (Glasser et al., 2013; Van Essen et al., 2013). The emotion task (176 volumes) was removed in the analyses because its minimal amount of volumes did not meet the low-motion criteria (details in the Study-2 section). For each task, there were two runs (one with LR phase coding direction and the other with RL). Thirty-one participants who have completed all the tasks and resting scans with low head-motion (mean FD < 0.25) were included in our analyses.

### HNU dataset

The Hangzhou Normal University (HNU) dataset is openly available via the Consortium of Reliability and Reproducibility (CoRR: http://fcon_1000.projects.nitrc.org/indi/CoRR/html/hnu_1.html) (Zuo et al., 2014). The dataset includes 300 scan sessions from 30 participants (15 males, age=24 +/- 2.41 years) who were scanned for 10 sessions over a month. Each session included an rfMRI scan acquired at the same GE MR750 3T scanner using a T2*weighted EPI sequence (TR=2000ms, TE=30ms, flip angle=90°, resolution=3.4×3.4×3.4mm, FOV=220mm2, 300 volumes, 10 mins). The details of this dataset were described in prior studies (Chen et al., 2015). We selected this dataset because the time duration between retest datasets is equally controlled across participants (3 days duration between two retest sessions) and the head motion was relatively low (mean framewise displacement [mean-FD] < 0.2mm for each of 300 scans, less than 1% [n=3] of time points within each scan has FD >0.2mm).

### MSC Dataset

The Midnight Scan Club (MSC) dataset consists of 10 participants, who each participated in 10 fMRI sessions over ten days beginning at midnight (https://openneuro.org/datasets/ds000224) (Gordon et al., 2017). Each session included 30 mins of resting fMRI using a gradient-echo EPI sequence (TR=2200ms, TE=27ms, flip angle=90°, resolution=4×4×4mm, 36 slices, 818 volumes). Consistent with the previous study, one participant with high head motion was excluded from the current work. The details of the imaging acquisition are described in the previous studies (Gordon et al., 2017; Gratton et al., 2018). Instead of using the temporal mask (FD<0.2mm) provided with the preprocessed data, we opted for a slightly less conservative threshold (FD<0.3mm) for further censoring high-motion time points to control the head motion while ensuring enough time points remained per scan (> 300TR).

### Data Preprocessing

#### HCP and HCP test-retest datasets

We used the publicly available version of the dataset that has been processed with the HCP minimal preprocessing pipeline (Glasser et al., 2013). In line with prior resting-state studies, nuisance regression, and bandpass filtering (0.01-0.1Hz) were then applied (Elliott et al., 2019). The nuisance regressors included Friston’s 24 head motion parameters, mean signals from white matter (WM) and cerebrospinal fluid (CSF), and linear and quadratic trends. The impact of regression with and without global signal was tested in our analyses. Given our focus on functional connectivity, nuisance regression, and bandpass filtering (0.01-0.1Hz) were carried out for minimal preprocessed data from task conditions. In addition, we examined the impact of task-evoked signals on our findings by adding regressors to account for task events/stimuli (Fair et al., 2007). Finally, the preprocessed data were projected to the cortical surface and smoothed with FWHM=6mm along the surface.

#### MSC

We made use of the preprocessed version of the MSC that is publicly available on OpenfMRI (https://www.openfmri.org/dataset/ds000224/) (Gordon et al., 2017; Gratton et al., 2018). The preprocessing steps included for this data included slice timing correction, grand mean scaling, and motion correction. The fMRI data were co-registered to the first session for distortion correction followed by detrending, nuisance regression (Friston’s 24 motion parameters, WM, CSF, and global signal) and bandpass filtering (0.009-0.08Hz). The details of the preprocessing steps were described in Gordon et al., 2017.

#### HNU

The HNU dataset was preprocessed using the Connectome Computation System (CCS: http://lfcd.psych.ac.cn/ccs.html) (Xu et al., 2015), a functional connectivity-focused end-to-end analytic pipeline. Briefly, preprocessing steps for functional data using the CCS included: removal of the first 5 TR’s, despiking, motion correction, grand mean scaling, Friston 24-parameter model nuisance regression and bandpass filtering (0.01-0.1Hz). The preprocessed fMRI data were co-registered to anatomical image and projected to the native midthickness surface and smoothed with FWHM=6mm along the surface.

### Functional Connectivity and Networks

To calculate functional connectivity matrices for each participant, vertex-wise time-series were first averaged within each of the 360 regions using a recent multi-modal parcellation atlas (Glasser et al., 2016). Pairwise Pearson correlations were calculated to generate a 360 x 360 parcel-to-parcel connectivity matrix. Reliability was calculated for each functional connection in the upper triangle of connectivity matrices (64620 connectivities). Yeo-Krienen 7 Networks was used to cluster the parcel-wise connectivity into large-scale networks (visual, somatomotor, dorsal, and ventral attention, limbic, frontoparietal, and default) (Yeo et al., 2011). The network identity was assigned to each parcel using the winner-take-all approach based on parcel-to-network overlap (i.e. percentage of parcels covered by the network). In addition, we also measured the functional connectivity using the partial correlation (Smith et al., 2013, 2011). Since the efficient calculation of partial correlation involves shrinkage estimation, the optimal regularization parameters might be subject- and state-specific. Here, we employed the partial correlation without regularization (Matlab function partialcorr). The Schaefer parcellation with a relatively smaller number of parcels (N=100, (Schaefer et al., 2018) was selected to reduce the computation costs. We also calculated the full correlation based on the Schaefer100 parcellation to ensure the robustness of the current study. In the following section, we focused on the full correlation results without specification, and reported the partial correlation in the supplement.

### Test-retest reliability Analyses

#### Terminology

To avoid a possible misunderstanding of terms used in the following section, we first clarify four key terms here: 1) *Scan* – a contiguous unit of fMRI data, defined by the start and stop of the MRI scanner. 2) *Session* – a unit of time that contains either a single or series of MRI experiments, defined by the participant entering and exiting the MRI scanner. 3) *Segment* – an epoch of contiguous fMRI time-series that is cut from a scan for data concatenation over more scans or sessions. 4) *Subset* – in the current study, since the test-retest dataset for reliability calculation is not necessarily acquired from a single scan or session, we define each of the test and retest datasets as a subset regardless of whether the data were from a contiguous scan or concatenated from multiple scans. As commonly performed in the literature, z-score normalization was conducted for each of the segments before concatenation.

#### Intra-class correlation (ICC)

Intra-class correlation for functional connectivity was calculated to estimate test-retest reliability using a Linear Mixed Model (LMM) (Chen et al., 2015; O’Connor et al., 2017; Xu et al., 2016; Zuo et al., 2013). For a given connection *c* denoted as *Y_ij_*(*c*), *i* represents the participant and *j* indicates the test and retest subset. Age, gender, global mean for functional connectivity, and head motion were included in the model as covariates.

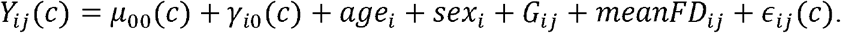

The intercept *μ*_00_(*c*) in the model represents the fixed effect of group average at connectivity *c*, while *γ*_*i*0_(*c*) represents the random effect for *i*th participant. Age, sex, and mean FD were included as the covariates in the model. In addition, as suggested in the prior study, we also included the global mean of functional connectivity (*G_ij_*) in the model to standardize the raw connectivity estimates *Y_ij_*(*c*) for *i*th participant and *j*th subset (Yan et al., 2013). The inter-individual variance 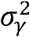 of the random effect and the variance 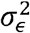 of residual *∈_ij_* were estimated using the restricted maximum likelihood (ReML) method in R (http://cran.r-project.org/web/packages/nlme). ICC was calculated by dividing the inter-individual variation 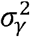 by the sum of 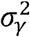 and residual variance 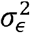. The ICC values are categorized into four intervals: poor (0< ICC≤0.4), moderate (0.4<ICC≤0.6), good (0.6<ICC≤0.8) and excellent (0.8<ICC≤ 1) in the results (Landis et al., 1977; Zuo and Xing, 2014).

#### General study flow

In this study, we examined the impacts of data aggregation on the reliability of functional connectivity from different data types (i.e. resting, task, hybrid), and concatenation methods. Figure 1 illustrates the specific comparisons of data aggregation strategies. More detailed descriptions for each study are introduced in the following sections.

**Figure 1.**
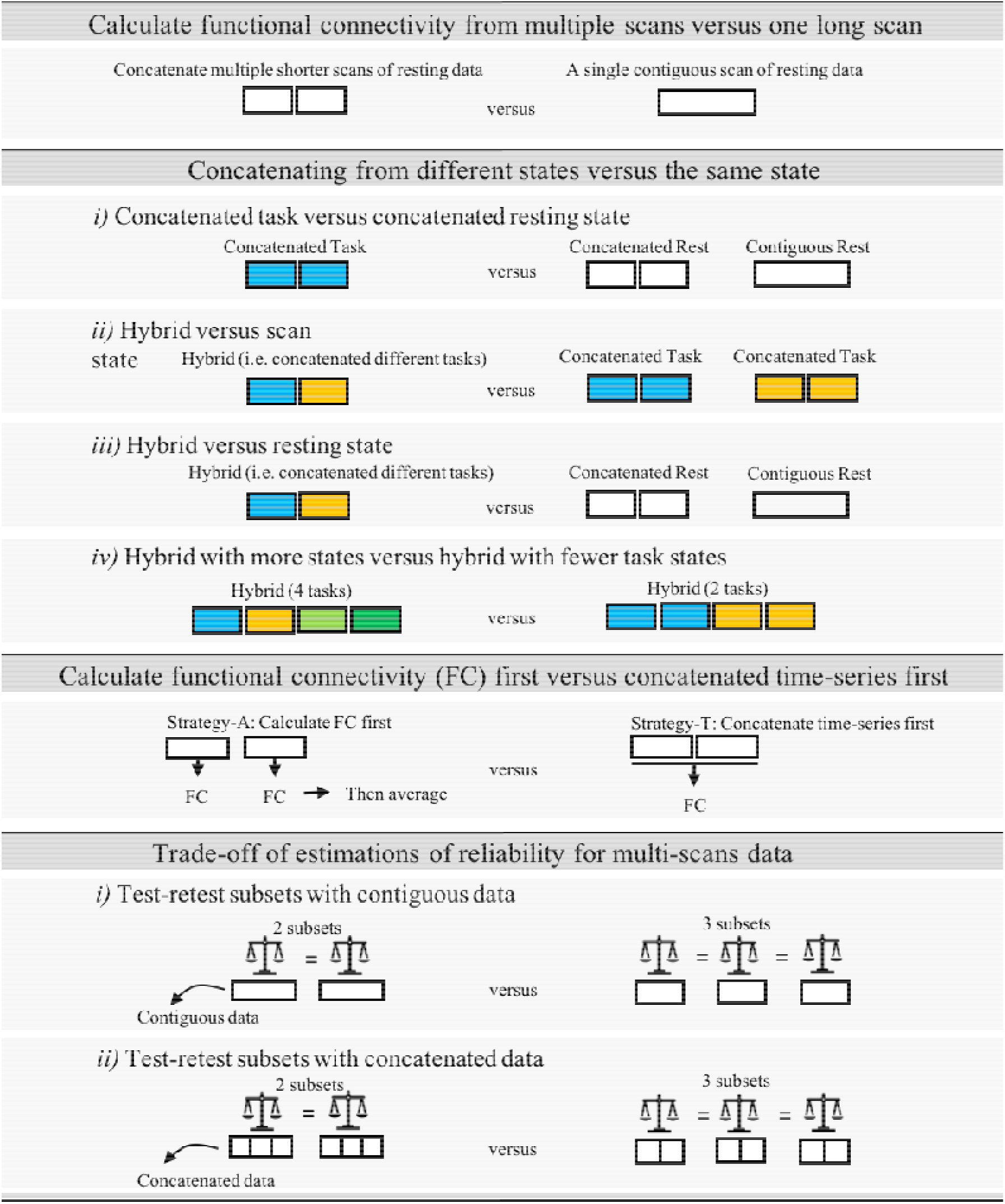
Graphical summary of comparisons for different data aggregation strategies.

### Study-1. Concatenating from multiple scans versus a single long scan

In order to address whether the protocol that required multiple shorter resting scans rather than a single scan could achieve comparable or higher reliable functional connectomics, we first examined the reliability of concatenating data from multiple scans versus that from a single long scan. Three datasets with multiple scans separated by a regular time interval between retest sessions were used in this analysis: 1) the HCP (n=217), which included 2 sessions per participant (2 R-fMRI scans per session; 1 day apart between two sessions), 2) the HNU (n=30), which included 10 R-fMRI scans per participant (3 days apart between two sequent sessions), and 3) the MSC (n=9) with 10 R-fMRI scans per participant (1 day apart between two sequent sessions). Of note, to maintain the continuity of R-fMRI scans, we performed the above analyses on the uncensored dataset and replicated this in the low-motion subsample of HCP dataset as well.

For the HCP dataset, we calculated the reliability of contiguous data across sessions (i.e. between Day 1 and Day 2), specifically, between Day 1 LR and Day 2 LR scans, and between the Day 1 RL and Day 2 RL. For the concatenated data, we used two resting fMRI scans (LR and RL, 600 TRs per scan) collected in Day 1 as the data pool to extract time-series segments and concatenate into subset 1. Similarly, data segments selected from two scans (LR and RL) in Day 2 were concatenated into subset 2.

Of note, the used scans matched on sequence (i.e. only LR or only RL) for contiguous data represents the test and retest of 15 mins scans (1200 TRs) with the same phase coding sequence for calculation of test-retest to obtain the best-case scenario for the reliability of contiguous data. An obvious concern that can arise is that the concatenated data combines across phase-encoding directions, while the contiguous data did not. To address this concern, we also calculated reliability for the scenario where they are not matched with the sequence to estimate the worst case direction over a day (Fig. S1A).

In the HNU dataset that consists of 10 sessions per participant, sequential sessions were used as test-retest subsets (i.e. test-retest between session #1 and #2, #2 and #3,…, #9 and #10) to examine the reliability of contiguous scans (280 TRs, ~ 10 mins). For the concatenated data, we first split 10 sessions into two data pools to generate test and retest subsets (Fig S1). Specifically, we used the first five scans (i.e. sessions 1-5) as the data pool to generate concatenated data for subset 1 and the last five scans (i.e. sessions 6-10) for subset 2. Similarly, different combinations of numbers of segments and lengths of segments to generate concatenated data were applied (i.e. 2 segments x 140 TRs, 4 segments x 70 TRs, 8 segments x 35 TRs). For the concatenated subset with 2 or 4 segments, each segment was randomly selected from a single scan to guarantee the concatenated data were merged from different scans. To ensure no contiguous segments in a single scan were selected in creating a concatenated subset with 8 segments, we randomly selected four scans from the data pool and then randomly selected segments from half of each selected scan to generate 8 segments into a subset.

In the MSC dataset, which contains 10 scan sessions per participant, a procedure similar to that applied in the HNU dataset was carried out to yield test-retest subsets (Fig S1). In short, the reliability was examined for contiguous data between two sequential sessions (800 TRs, ~ 30-mins per session), as well as concatenated data from different combinations (i.e. 2 segments x 400 TRs, 4 segments x 200 TRs, 8 segments x 100 TRs, 16 segments x 50 TRs, 800TRs [30 min] in total per subset).

### Study-2. Concatenating from different states versus the same state

A recent study suggested that functional connectivity calculated by concatenating multiple fMRI task scans in the HCP dataset exhibits higher test-retest reliability than that from an equivalent amount of pure resting-state fMRI data (Elliott et al., 2019). While this may be true, prior work using the MyConnectome dataset highlighted the advantage of concatenation from shorter scans over a single scan (Laumann et al., 2015). As such, it is not clear whether the better reliability for merged data from different states (referred to here as the ‘hybrid’ condition) is driven by the concatenation advantage or the combination of experiment states. To address this question, we generated the concatenated resting subsets that were matched to the hybrid condition with respect to the number/length of segments and compared the reliability between hybrid and concatenated resting data. In addition, we further examined the impact of concatenation from combinations of states.

#### Concatenated task versus concatenated resting state

We first evaluated the reliability of concatenated data over available scans (2 scans, LR and RL, 5.6 min in total) from the same task only versus resting state (2 scans, LR and RL, 5.6 min in total). We also compared the reliability of hybrid data yielded from two different fMRI tasks as compared to the concatenated data from a single state (task or resting). The reliability of six task states, resting state, as well as the hybrid from two different tasks were calculated. The effects of global signal regression (GSR) and task regression were examined in our analyses. Of note, we used the resting without ICA-FIX, consistent with the preprocessing procedure of task data. We also tested the ICA-FIX resting data in the supplementary materials (Fig. 7, S6).

#### Hybrid versus concatenating from resting state

We made use of the HCP test-retest dataset to assess the reliability of hybrid versus resting-state conditions between test and retest measurements. Specifically, we used six tasks to generate all of the possible hybrid subset combinations with regard to task state and encoding direction (LR and RL), consisting of 2 different tasks (2 tasks x 232 volumes, 5.6 min), 2 different tasks that each contains 2 scans (2 tasks x 2 scans x 232 volumes, 11.1min), 4 different tasks (4 tasks x 232 volumes, 11.1min) and 6 different tasks (6 tasks x 200 volumes, 15 min). To match the number of segments of hybrid data, concatenated resting-state subsets for comparisons were created by concatenating 2, 4, and 6 segments. The segments were contiguous volumes (i.e., the first 232 or 200 volumes) from different resting scans, except for the 6-segments combination in which two additional segments were selected (#601-800 volumes) since only 4 resting scans were acquired. In addition, we compared the reliability of concatenated hybrid and resting connectivity against that from a single contiguous resting scan truncated to the same time duration.

### Study-3. Calculate and then combine versus Combine and then Calculate

When multiple scans are acquired, apart from combining time-series first, we can also calculate the functional connectivity per scan then average the connectivity afterwards. In this section, we have tested whether one strategy is superior to the other using the HCP and HNU datasets.

In the HCP dataset, resting fMRI data from Day 1 LR and RL scans were used for 1 subset while Day 2 LR and RL scans were used for the 2nd subset. Reliability was tested between the Day 1 subset and Day 2 subset. The functional connectivity matrix for each subset was either calculated from concatenated data over LR and RL scans, or averaged from connectivity that calculated from each of LR and RL scans separately. We tested 3 different time lengths, 1) 5 minutes by 2 scans per subset, 2) 10 minutes by 2 scans per subset, and 3) 15 minutes by 2 scans per subset.

In the HNU dataset, we selected two scans (session #1-2) to create subset 1 while two scans (session #6-7) for subset 2. In this way, the time duration between test-retest subsets are fixed to 12 days (scanned once every 3 days). Functional connectivity was calculated in both strategies from two sessions per subset. Similarly, we tested 2 different time lengths, 1) 5 minutes by 2 sessions per subset, and 2) 10 minutes by 2 sessions per subset.

### Study-4. Delineating the trade-off of amount of data per subset and number of subsets in evaluations of reliability

Beyond the general rule of ‘the more the better’ in data collection, in this section, we focused on how test-retest reliability increases with more numbers of subsets versus more amount of data per subset. We examined the reliability for different combinations of these two factors.

#### Test-retest subsets with concatenation

The MSC dataset was used here as it contains 10 sessions x 30 mins data per subject. We tested the number of subsets varied from 2, 3, 4, 5, and 6; the amount of time per subset varied from 10, 12.5, 15, 17.5, 20, 22.5, 25, 30, 35, 40, 50, and 60 mins. For a combination of *k* subsets and *t minutes* per subset, the subset data was concatenated from *x* numbers of short contiguous segments (2.5 mins per segment, *x*=*t*/2.5). Specifically, for each participant, we first randomly cut all available data into a few 2.5 mins (68 volumes) segments as a segment pool. We then randomly selected *k*x* segments to create *k* subsets. The test-retest reliability was calculated for all combinations of numbers of subsets and the amount of data per subset. We replicated the analyses in the HCP dataset for different combinations of 2, 3, 4, 5 subsets by 10, 12.5, 15, 17.5, 20, 22.5, 25 mins of data per subset. The subsets were concatenated in a similar way from short segments (2.5 mins, 208 volumes).

#### Test-retest subsets without concatenation

We also considered the test-retest subset acquired naturally in a single fMRI session instead of concatenated from multiple shorter scans in the MSC dataset. Similarly, we tested the number of subsets varied from 2, 3, 4, 5, and 6, while the amount of time per subset varied from 10, 12.5, 15, 17.5, 20, 22.5, 25, and 30 mins. For a combination of *k* subsets and *t minutes* of scan duration, *k* sessions (*k*=2,3…,6) were randomly selected *k* sessions first and *t* min of contiguous time points from each of selected *k* sessions. The test-retest reliability was calculated for all combinations of numbers of sessions and scan duration.

## RESULTS

### Study-1. Concatenating from multiple scans versus one long single scan

Here, we compared the ICC distribution for edge-wise functional connectivities based on fMRI data concatenated from multiple scans, with that obtained from a single long scan. Generally, using the HCP data, we found that when the amount of data was equated, there was a clear advantage for multiple scans from the perspective of edgewise reliability. Specifically, the concatenated subset (7.5 min from each of 2 scans) yielded higher ICC’s than the contiguous data (15 min from a single scan) (Fig. 2A, ICC matrices in Fig S2B). To test the significance of the ICC improvement for concatenated data, we performed the permutation test by randomly shuffling the labels (concatenated vs. contiguous) 10,000 times. We found that significance did not depend on whether GSR was included in preprocessing or not (without GSR: *p*(*permute*) <10^-4^, effect size = 0.425 [0.436-0.414], GSR: *p*(*permute*)<10^-4^, effect size = 0.209 [0.198,0.220]). Such advantages of concatenation were observed across all networks (Fig. 2B). Similar findings were obtained in the HNU (contiguous scan duration = 10 minutes, Fig. 2C-D) and MSC datasets (contiguous scan duration = 30 minutes, Fig. 2E-F), as well as the low-motion HCP subsample--all achieving statistical significance (Fig. S3A).

**Figure 2.**
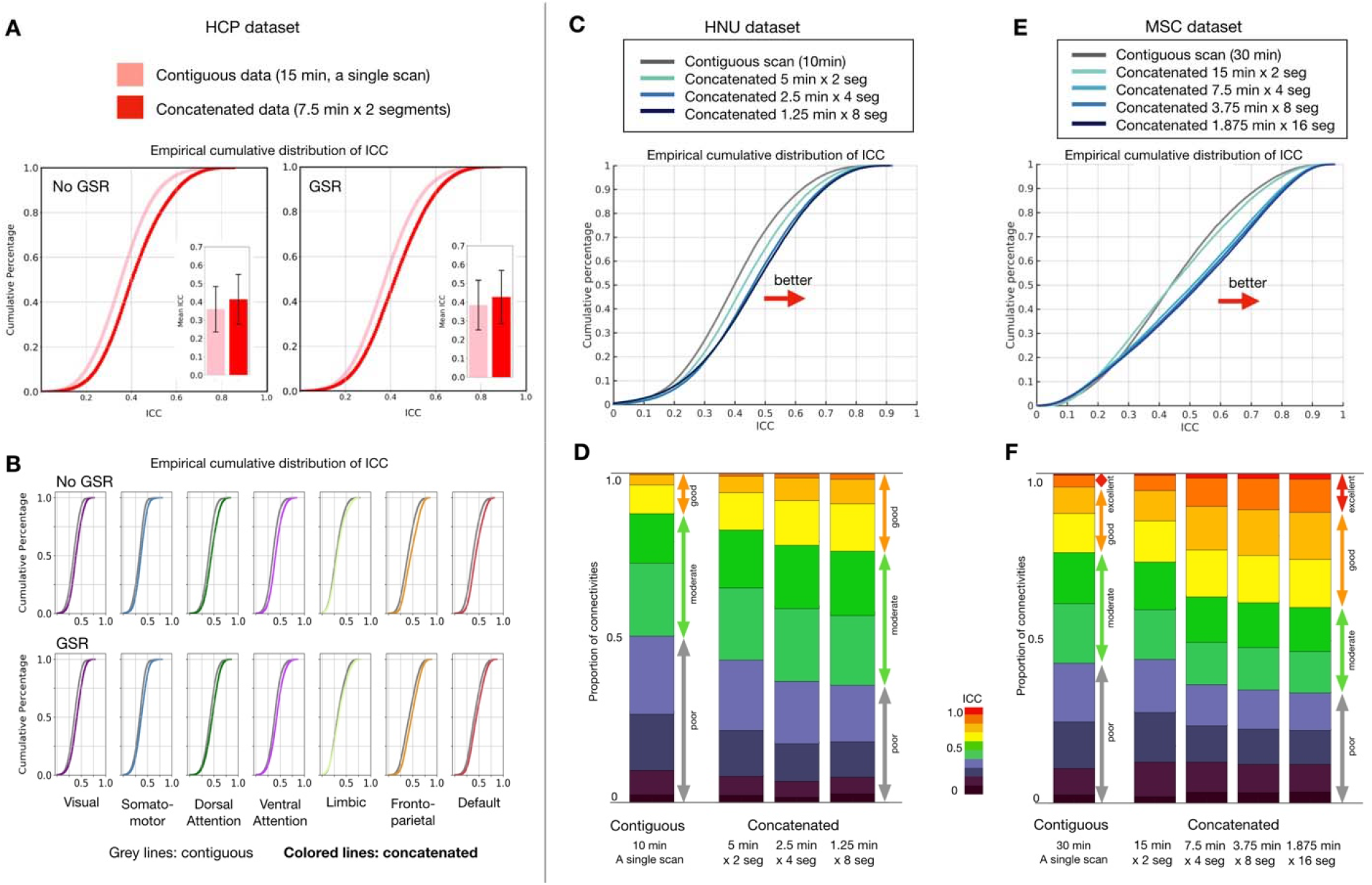
Edgewise functional connectivity calculated from concatenated scans is more reliable than that based on a single contiguous scan of equal length. **(A)** Empirical cumulative distribution and bar plots of mean intraclass correlation coefficient (ICC) with error bars (standard deviation) for test-retest reliability of functional connectivities from contiguous and concatenated resting-state fMRI scans; results for calculation based on data with and without global signal regression (GSR) are depicted. **(B)** Empirical cumulative distribution of ICC for each network. Distribution of ICC from contiguous data is shown in grey while colored lines represent ICC from concatenated data. See FC and ICC matrices in Fig S1B. **(C)** Empirical cumulative distribution of ICC and **(D)** stacked bar plots depicting proportion of connections with poor, moderate, good and excellent ICC for contiguous and concatenated resting-state scans, calculated using the HNU dataset and the Midnight Scan Club dataset **(E, F)**.

With regard to the concatenation window size, we found that from a reliability perspective, a higher number of shorter fMRI data segments appeared to engender advantages over a smaller number of larger segments. In the HNU dataset, 10 min of a contiguous scan yielded 12% of connections with good reliability (ICC>=0.6), which increased to 26% for 8 x 1.25 min segments from four sessions. In the MSC dataset with 30 min of data, shorter segments consistently showed higher ICC than contiguous data (contiguous: 24% of connectivities were good; 16 x 1.875 min segments: 41% good). Of note, this advantage does not appear to result from the process of Z-scoring (i.e., mean centering and variance normalization) segments prior to combining; there was no advantage, and even slightly worse reliability, observed when segments were selected from the same scan (Fig. S2C). We also tested whether concatenating time-series data from the onset of each scan could achieve higher reliability, but did not find any clear evidence there is the onset effect (Fig. S3B).

We also examined the impact of the time duration in concatenated data. Several studies, including the present (Fig S2A), have demonstrated the data collected within a shorter duration (e.g. same day) were more reliable than those with longer duration (e.g. 1 day, 1 week or month apart) (Noble et al., 2019; Zuo et al., 2013). Therefore, we expected that the concatenated segments from a shorter duration (same day) would generate higher reliability. Here, we used the HCP dataset, and calculated the reliability of concatenated data from the back-to-back scans on the same day (rfMRI_REST1_LR, rfMRI_REST1_RL, 7.5 min from each of 2 scans) as compared to concatenated data from two different days (i.e. 1 day apart, rfMRI_REST1_LR, rfMRI_REST2_LR, 7.5 min from each of 2 scans). As we expected, the segments selected from scans with a shorter duration on the same day yielded higher reliability than those over two days (Fig S3C).

In addition, we also tested the concatenation effect for partial correlation using the HCP data. In addition to ICC, we also calculated the discriminability to examine the reliability for the whole cortical connectome (Bridgeford et al., n.d.). Consistent with previous studies, we found that the reliability of the partial correlation was lower than the full correlation for both contiguous and concatenated data (Table S6). However, when comparing the reliability of data from multiple scans versus a single scan, the partial correlation also showed an advantage in concatenated data over contiguous data (permutation *p*<10^-4^).

### Study-2. Concatenating from different states versus the same state

Study-1 showed that the concatenation of data from multiple scans yielded connectivity estimates with higher reliability than those obtained using a single scan. However, the data was all obtained during the resting state. Here, we take the next step, and examine: 1) the reliability of concatenated data from a task state versus resting state, 2) the reliability of concatenated data from different states (rest, task) versus the same state, 3) whether the benefits of concatenation are maintained when concatenating data from multiple states (i.e. hybrid) versus a single state, and 4) the factors that potentially contribute to the reliability of hybrid concatenations. We focus on the full correlation in the following sections and partial correlation results in the supplement (Fig. S9).

### i) Concatenated task versus concatenated resting state

Using HCP test-retest dataset, we concatenated the available task scans (232 volumes x 2 scans, 5.6min in total) for each task state and compared their reliabilities to the concatenated resting data (232 volumes x 2 scans), as well as the contiguous resting data (464 volumes from a single scan). We found that when task regression was performed, edgewise functional connectivities for two of the task states (i.e., language and social) exhibited higher ICC’s than those from either concatenated or contiguous resting data. When task regression was not applied, reliability scores for edgewise connectivities increased for all six task states, becoming higher than those from resting state (Fig. 3A). Similar findings were observed when GSR was carried out (Fig. S4).

**Figure 3.**
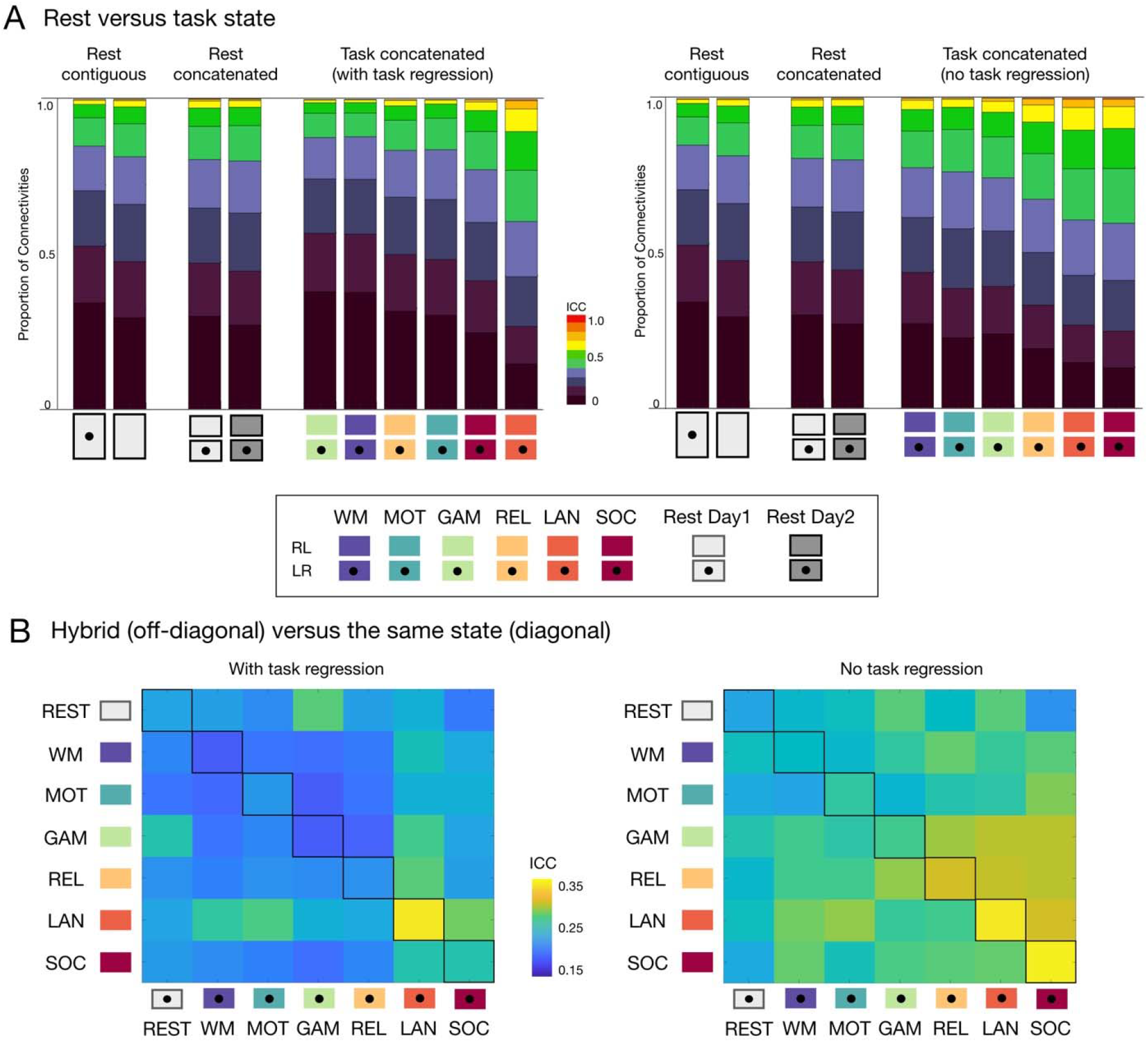
Reliability of edgewise functional connectivity measures calculated for resting-state, task and hybrid fMRI data without GSR. **(A)** Proportion of connections with poor, moderate, good and excellent ICC when calculated for resting state and each of six task conditions in the HCP dataset, with (left-panel) and without task regression (right-panel). **(B)** Mean ICC for functional connectivity measures based on hybrid data generated by concatenating two scans from two different states (off-diagonal, upper triangle: LR scans, lower triangle: RL scans) and the same state (diagonal). See results with GSR in Fig. S4.

### ii) Hybrid versus the same state

Next, we considered data concatenated from 2 different states, instead of 2 scans from the same state. We systematically tested all the possible combinations of concatenating 2 different states. The concatenated language task with task regression showed the highest reliability among the possible combinations of different states containing the language task (Fig. 3B, left). When task-regression was performed (Fig. 3B, right), three concatenated tasks (language, social, and relational) showed higher reliabilities than their hybrid combinations (off-diagonals). It is important to note that hybrid data generated from two states associated with more reliable states (e.g., language and social tasks) tended to yield greater reliabilities than the other possibilities (e.g., motor and working memory tasks).

### iii) Hybrid versus resting state

We further compared the reliability of edgewise connectivities from multiple fMRI scan types (i.e. hybrid) to those from resting fMRI data, concatenated from multiple scans, as well as from equal amounts of resting data from a single scan. Besides the combination of 2 states (2 scans, 5.6 min in total) that were tested above, we also tested the hybrid connectivity of 2 states generated from a higher number of scan types (4 scans, 11.1 min in total). The concatenated resting data (equal length) were generated from 2 and 4 resting scans respectively, as well as equivalent length of contiguous resting data truncated from a single scan (Fig. 4). As expected, more data was better, regardless of hybrid or resting state. Concatenated resting data (blue dot) exhibited higher reliabilities than half of the 21 possible hybrid combinations from 2 scans with task regression. When compared with hybrid from 4 scans, the concatenated resting data yielded relatively higher reliabilities than most of the hybrid combinations (17 out of 21 combinations), except for the hybrid generated among language, social and rest (Fig. 4). In contrast, when task regression was not performed, the hybrid data showed generally higher reliabilities than concatenated and contiguous resting data (Fig. 4, right) combinations. Similar trends were found for GSR preprocessed data as well (Fig. S5).

**Figure 4.**
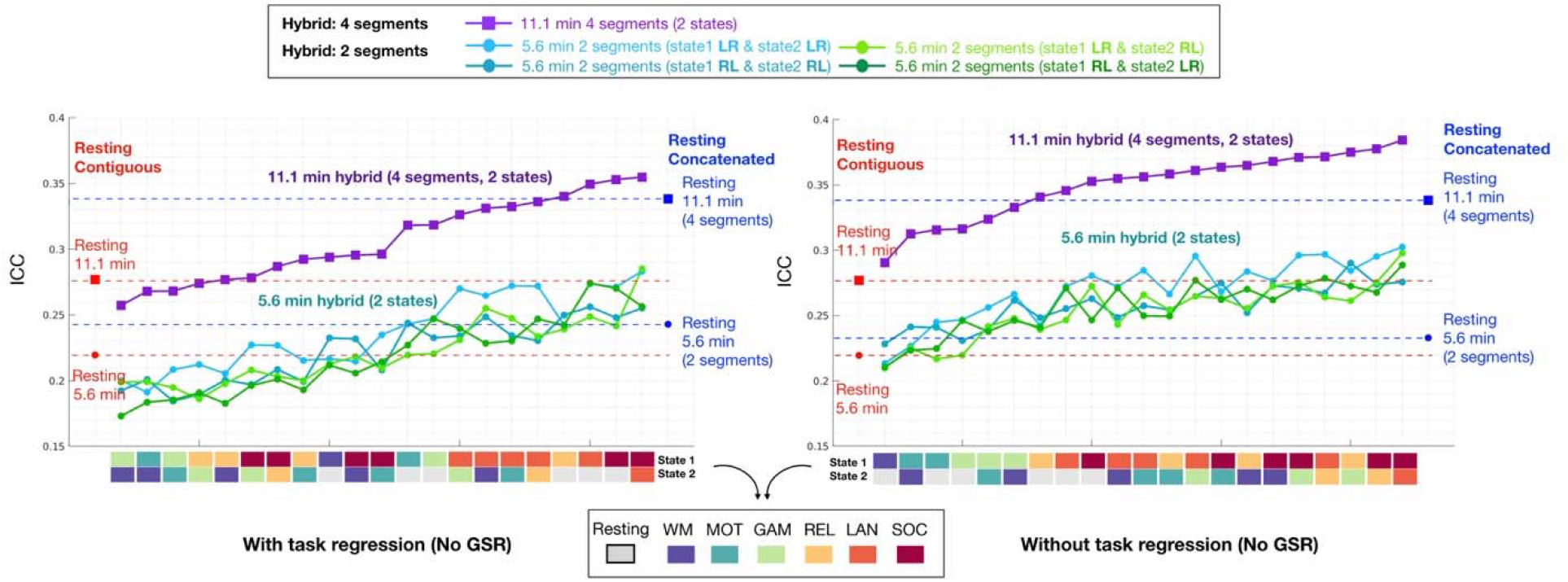
Reliability of edgewise functional connectivity measures calculated for resting and hybrid data generated from 2 tasks using either 2 or 4 segments without GSR. The label color for task states from cold to warm represents the reliability (i.e., mean ICC) of each state from low to high (detailed distribution shown in Fig. 3A). The same length of contiguous resting data was truncated from LR scans of the Day 1 session (red dots, 5.6 min; red square: 11.1 min). The concatenated resting data was generated from 2 scans (blue dot: 5.6 min, LR and RL scans from Day 1) and 4 scans (blue square: 11.1 min, LR, RL scans from Day 1 and Day 2). See results with GSR in Fig. S5.

Besides the combinations of 2 states we tested above, we also used 4 and 6 task states to generate different lengths of hybrid data, and systematically tested all possible combinations of different tasks (details in Method). We found that compared with contiguous resting data, that hybrid data generated from 2 tasks (total data: 5.6 min), 4 tasks (11.1 min), or 6 tasks (15 min) did not necessarily exhibit higher reliabilities (Fig. 5). When task-evoked activation was regressed out (Fig. 5A), nine out of forty-two combinations of the shorter, two-states hybrid had an advantage over the concatenated resting-state with respect to reliability (Fig. 5, left panel). With more data (hybrid: 4 or 6 different tasks, resting: 4 or 6 segments), the concatenated resting data showed a clear advantage over all hybrid conditions and contiguous resting as well (Fig. 5, middle and right panels). Similar findings were observed in data with GSR (Fig. 5A, gray line). When matching the number of concatenation segments for resting conditions, the concatenated resting data offered higher reliability than most of the hybrid combinations from different task states (Fig. 5A, gray triangles on the blue lines).

**Figure 5.**
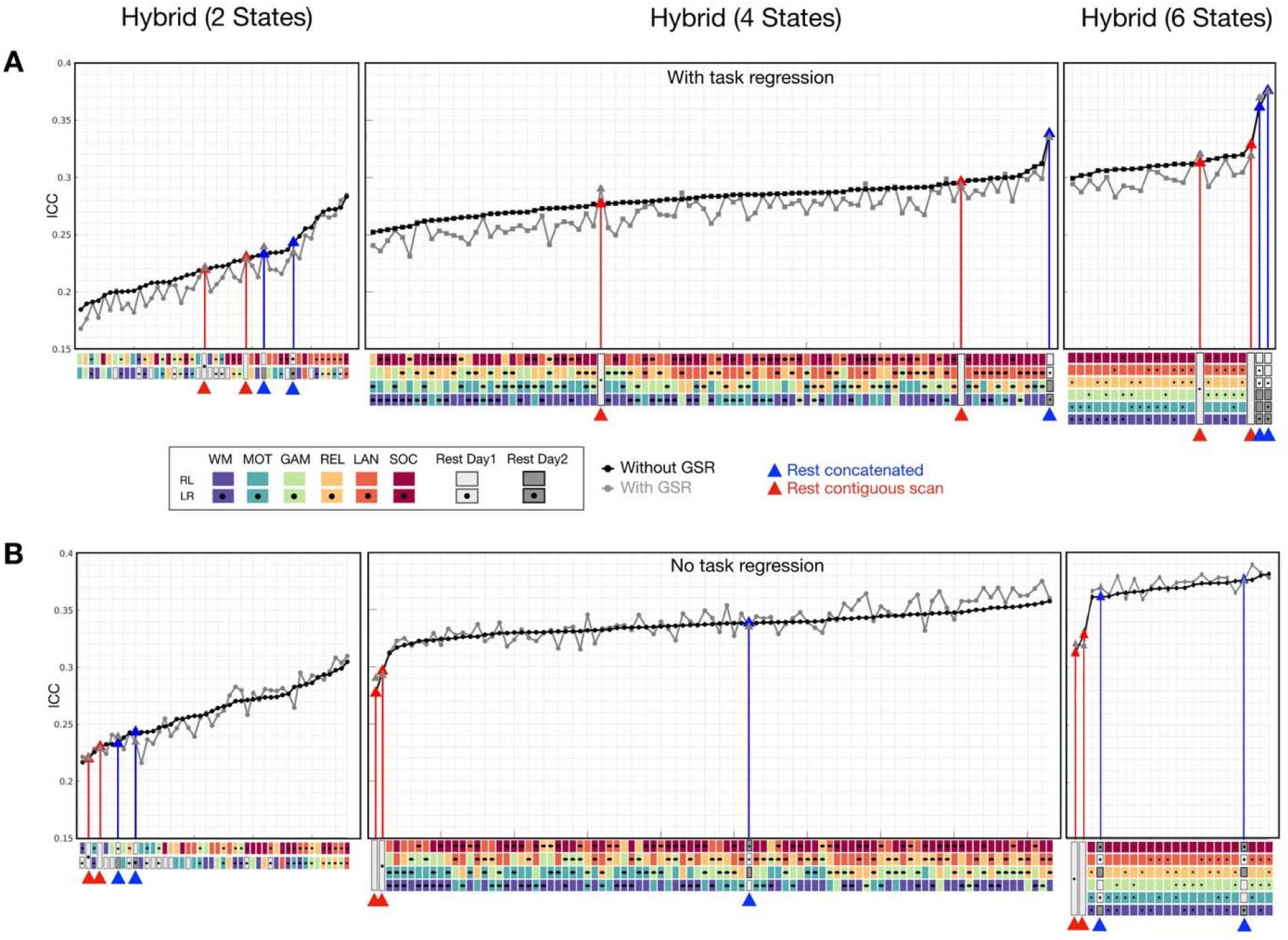
Reliability of edgewise functional connectivity for resting and hybrid data as a function of the numbers of states combined in the concatenation process (2, 4 and 6 different states). The mean ICC for data preprocessed with GSR (gray) and without GSR (black) was shown in ascending order. The same length of contiguous resting data was truncated from LR and RL scans of Day 1 session (highlighted with red lines). The concatenated resting data was generated from 2, 4, and 6 segments of resting scans (highlighted with blue lines).

We also assessed the ICC of hybrid conditions from task scans without task regression. As depicted in Fig. 5B, connectivity reliabilities for hybrid from task states were higher without task regression than with. Focusing on the results without GSR first (black line), when comparing such hybrid conditions with contiguous resting data, the hybrid conditions consistently yielded higher ICC’s than contiguous resting data (red triangles). The concatenated resting data (blue triangles), on the other hand, showed increased ICC and became more likely to have higher reliabilities than hybrid combinations from more than 2 states (i.e. 4 tasks and 6 tasks). A similar trend was observed with GSR data (Fig. 5B, gray line).

### iv) Hybrid with more task states versus hybrid with fewer task states

We previously noted that connectivity reliabilities from the resting state were not superior to those from all of the task states (Fig. 3A). It seemed that combining more different task states (>=4 tasks) might have a negative impact on reliability, resulting in lower reliabilities than resting state. Thus, we further controlled the number of segments and varied the number of task states combined. Specifically, we compared the reliabilities yielded from hybrid data concatenated from 4 different tasks (4 scans) as we examined above versus 2 different tasks (2 scans per task, 4 scans in total). As we expected (Fig. 6, dark green diamonds), hybrid combinations from fewer states (2 tasks) appeared to yield higher reliabilities than hybrid from 4 tasks. In particular, hybrid data generated from language and social tasks exhibited higher reliabilities than concatenated resting data. Of note, language and social tasks are also the most reliable tasks and showed higher reliability than resting state (Fig. 3A). These findings also suggest that data concatenated from reliable states tend to yield more reliable connectivities. Similar observations were found in data with GSR (Fig. 6, gray line, light green diamonds). The highest reliabilities were obtained for the hybrid data generated from fewer tasks, though less hybrid combinations were observed when task-regression was not performed.

**Figure 6.**
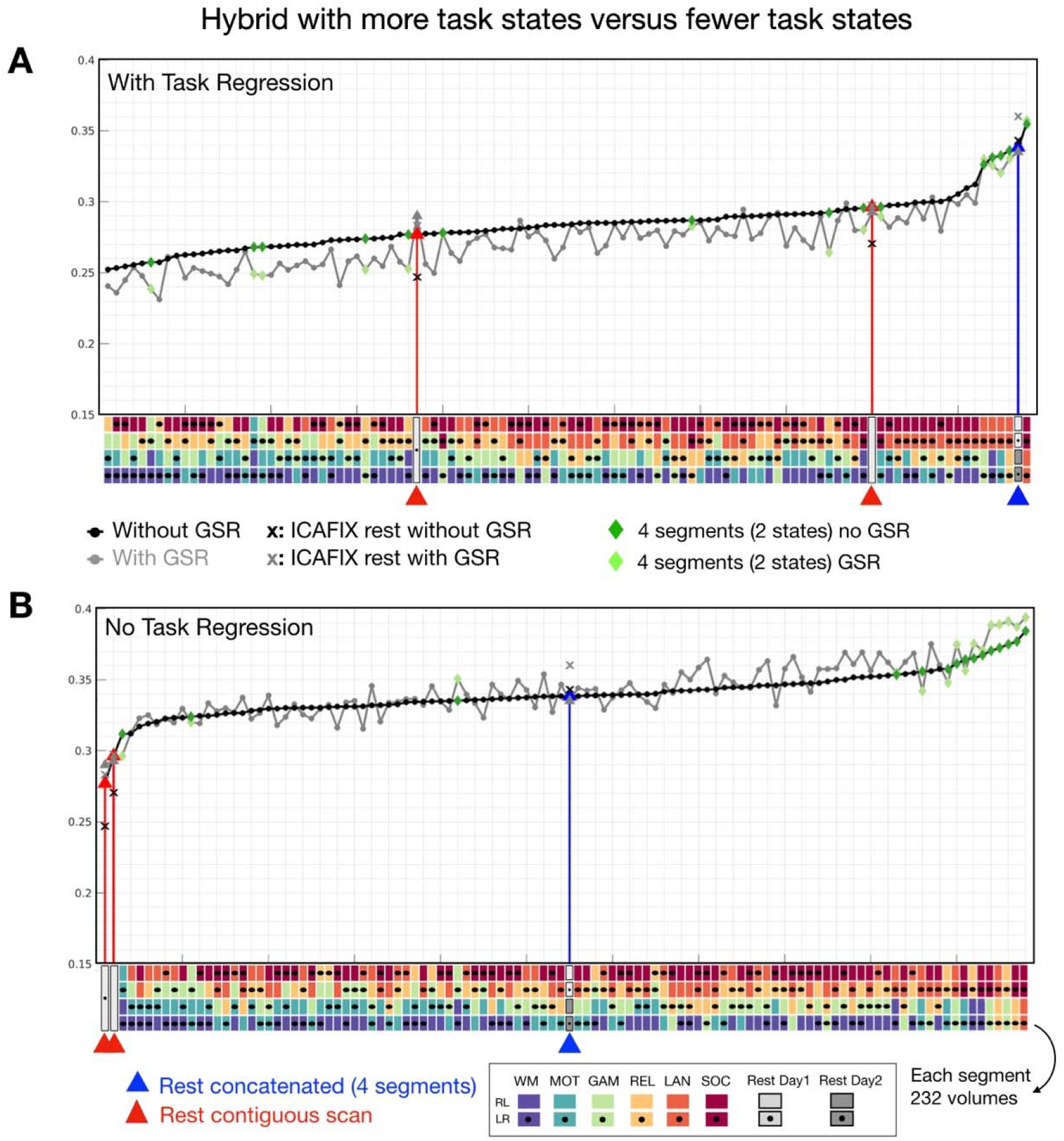
Reliability of edgewise functional connectivity for hybrid data as a function of the numbers of task states (2 versus 4 tasks) combined in the concatenation process. Hybrid combinations with less task states (green: 2 tasks) exhibited higher reliability than hybrid combining from more task states (black: 4 tasks).

#### v) What determines the reliability of hybrid data?

So far, our findings suggested that hybrid segments selected from more reliable states (e.g. language and social) and more homogeneous states (e.g. same state > different states) might obtain higher reliability. Thus, we further examined how these factors contribute to the reliability of hybrid data. Specifically, we evaluated 1) the averaged reliability of states used in hybrid concatenation and 2) the similarity of FC between states used in hybrid concatenation. As expected, we found that the reliability of the 2-segment hybrid data was significantly correlated with the mean reliability of the states used in concatenation (Fig 7A), while there was no relation with the similarity of FC between states (Fig 7B). Such correlations were consistent across all preprocessing pipelines (GSR and task regression). These findings suggest that hybrid data from 2 states were mainly driven by the reliability of state types used in the concatenation, not the homogeneity between states. However, when the hybrid data were concatenated from more states (i.e. 4 states), reliabilities of states did not exhibit any positive correlations with the reliability of the hybrid data (Fig 7C). Notably, a strong correlation was observed for the similarity of FC between states (Fig 7D). This suggests, when more state types were used in the hybrid concatenation, the homogeneity of the states, instead of the reliability of each state determines the reliability of the hybrid data.

**Figure 7.**
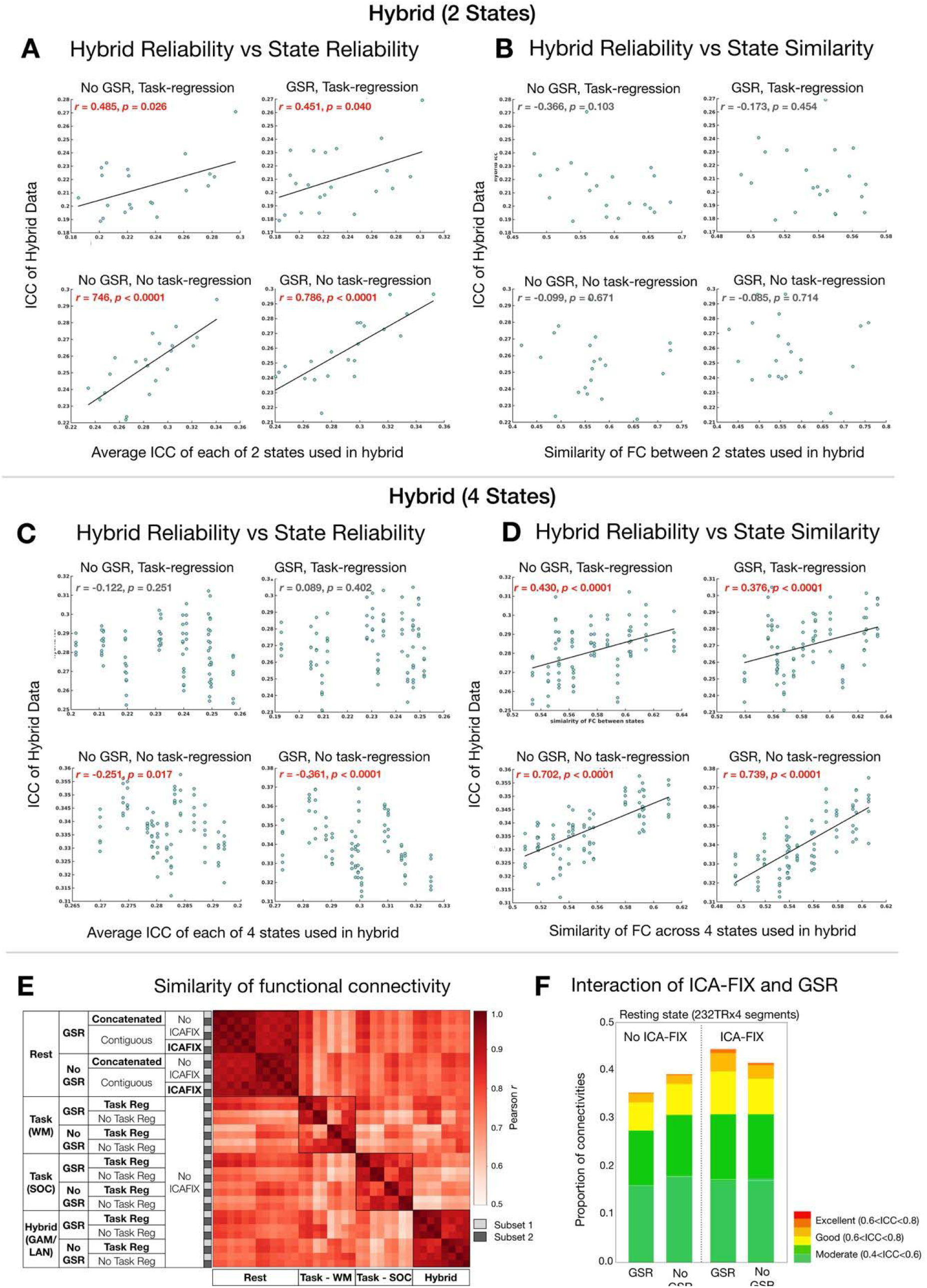
The factors that contribute to reliabilities of hybrid data and the effect of preprocessing steps for different states. **(A, C)** The correlations between the reliability of hybrid data and the average ICC of states used in 2-states hybrid (A) and 4-states hybrid (C). **(B, D)** The correlations between the reliability of hybrid data and the similarity of FC across states used in 2-states hybrid (B) and 4-states hybrid (D). **(E)** Averaged within-individual similarity (Pearson’s *r*) of functional connectivity patterns for rest, task, and hybrid data (5.6 min in total) with different preprocessing options. Working memory (WM) and social (SOC) state was shown as an example of the task condition and concatenation from GAM and LAN for the hybrid condition. The group averaged functional connectivity matrices with different preprocessing options are shown in Fig S6. **(F)** Proportion of connections with moderate, good, and excellent ICC for concatenated resting-state data preprocessed with and without both GSR and ICA-FIX.

We also noted that the shorter time duration yielded higher reliability for concatenated resting data (Fig S3C). Thus, we tested whether this true for hybrid data. Given that all HCP scans were collected in 2 days, we assigned the mean time duration to 1 if two states were collected on the same day otherwise 2. We then compared the reliabilities of hybrid conditions with a shorter and longer time duration. Unfortunately, we did not find significant differences with any preprocessing pipelines (GSR and task-regression). Similar non-significant findings were observed for hybrid data from four states. We suspect that the segment duration matters more when all segments are the same state (e.g. rest), whereas when concatenating different states, the segment duration is much less important than the reliability of states or homogeneity of FC between states, which showed the dominant factor determining the reliability of the hybrid data.

#### vi Dominant preprocessing steps for the task, resting and hybrid data

To examine the impacts of the preprocessing steps for different states and hybrid data, we first measured the similarity (Pearson’s *r*) of functional connectivity patterns within individuals under different preprocessing options across task, resting, and hybrid conditions. The group averaged functional connectivity with different preprocessing (i.e. GSR, ICA-FIX, task-regression) is shown in Fig. S6. We found that the GSR option dominated the functional connectivity patterns for resting, task as well as hybrid data (Fig. 7E). It is worth noting an interaction between ICA-FIX and GSR (Fig. 7F). Without ICA-FIX, no GSR option provided a higher ICC than GSR (permutation *p*<10^-4^), while GSR showed an advantage over no GSR when ICA-FIX was performed (permutation *p*<10^-4^). When compared to the hybrid data, the concatenated resting conditions with GSR and ICA-FIX exhibited higher ICC than most of the hybrid combinations (Fig. 6, “X” marks in grey color).

### Study-3 Calculate and then combine versus combine and then calculate

Besides time series concatenation, another data aggregation strategy is to first calculate the functional connectivity, then average connectivity matrices across scans. Here, we further compared these two strategies using the HCP dataset. We have included two scans, each scan varied from 5 min, 10 min, and 15 min for data combination. The strategy of concatenating the time-series first was denoted as Strategy-T, while calculating the FC per scan then averaging the FC was denoted as Strategy-A. We found no significant difference between Strategy-T and Strategy-A for 5 min, 10 min and 15 min data (Fig. 8). Similar findings were replicated in the HCP low-motion subsample and the HNU dataset (Fig. S7).

**Figure 8.**
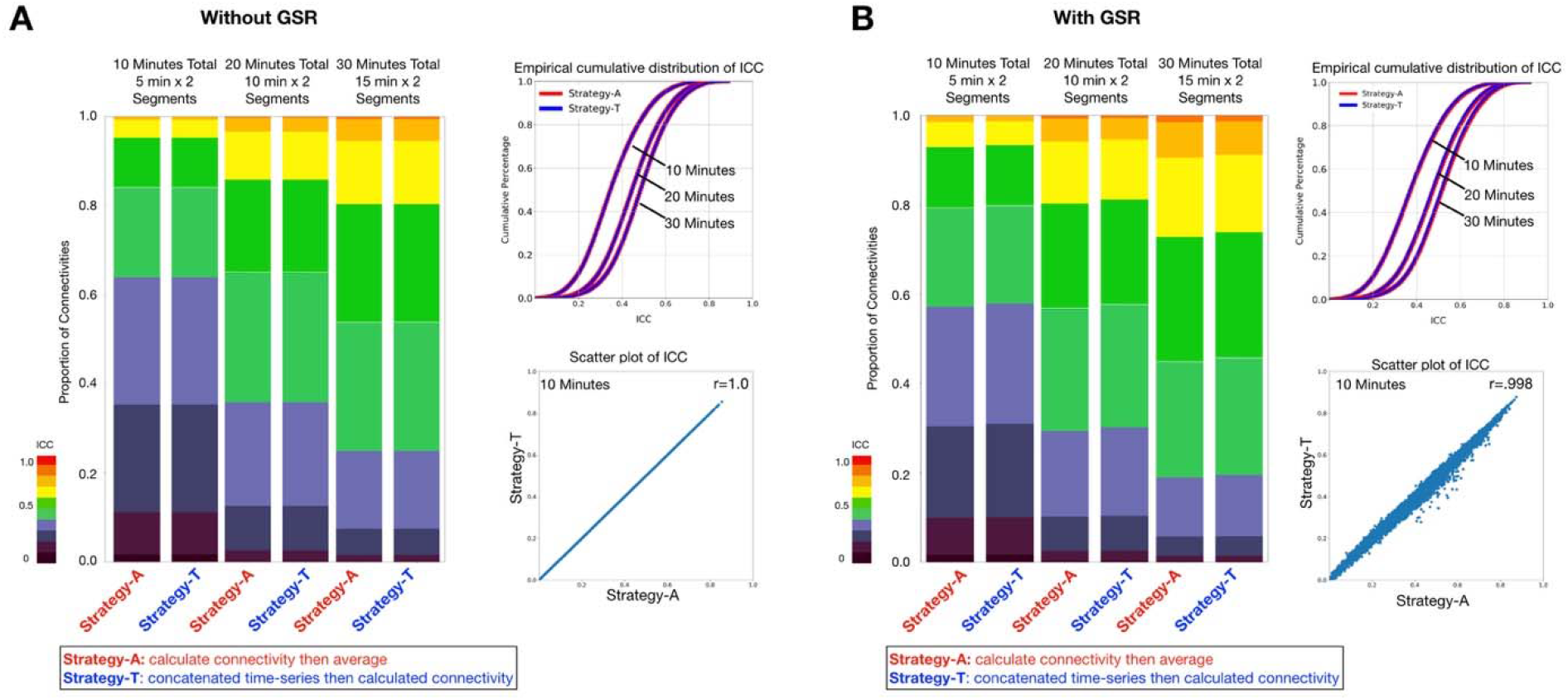
Calculation of edgewise functional connectivity measures for scans individually and then averaging, versus calculation after concatenation of scans. Two segments of 5 min, 10 min, and 15 min of data were generated. The strategy calculating the FC of each segment first then average all the FC as the final FC for each subset was denoted as Strategy-A, while concatenating time-series of all segments then calculating FC per subset as Strategy-T. The stacked bar and the empirical cumulative distribution of ICC showed no differences in ICC distribution between two strategies. The scatter plot shows the ICC of functional connectivity was almost the same between two strategies for 10 min data (2 segments x 5 min).

Intriguingly, when two strategies were compared using the Schaefer100 parcellation (Figure S10, Table S10), Strategy-T showed significantly higher reliability than Strategy-A for full correlation (*p*<10^-4^). Similar findings are observed for partial correlation when the subset time is 20 min (10 min per segment). However, the Strategy-A appeared to be superior to Strategy-T for data over 30 min (permutation *p*<10^-4^).

### Study-4. Delineating the trade-off of amount of data per subset and number of subsets in evaluations of reliability

In Study 1, we showed that more data per subset improved the reliability between two subsets, regardless of whether the subset was contiguous or concatenated data. These findings echoed the general rule of ‘the more the better’ in data collection. Here we address a confusion that may arise in evaluations of reliability--whether increasing the number of subsets improves reliability more quickly than increasing amounts of data per subset.

We first focused on the contiguous data using the MSC dataset (30 min x 10 sessions per subject). We calculated the test-retest reliability using 30 min in total, but estimated from two models 1) two test-retest subsets with 15 min per subset, and 2) three subsets with 10 min per subset. As shown in Fig. 9A, the reliability estimated from two subsets showed higher reliability than estimation from three subsets. We also tested the reliability for each subset created by concatenating smaller time-series segments (2.5 min) randomly selected within the subject from all available data without replacement (Fig. 9B). Similarly, when the total amount of the data was used (30 min), more data assigned to subsets (15 min x 2 subsets) yielded higher reliability than when the data was split into more subsets (10 min x 3 subsets).

**Figure 9.**
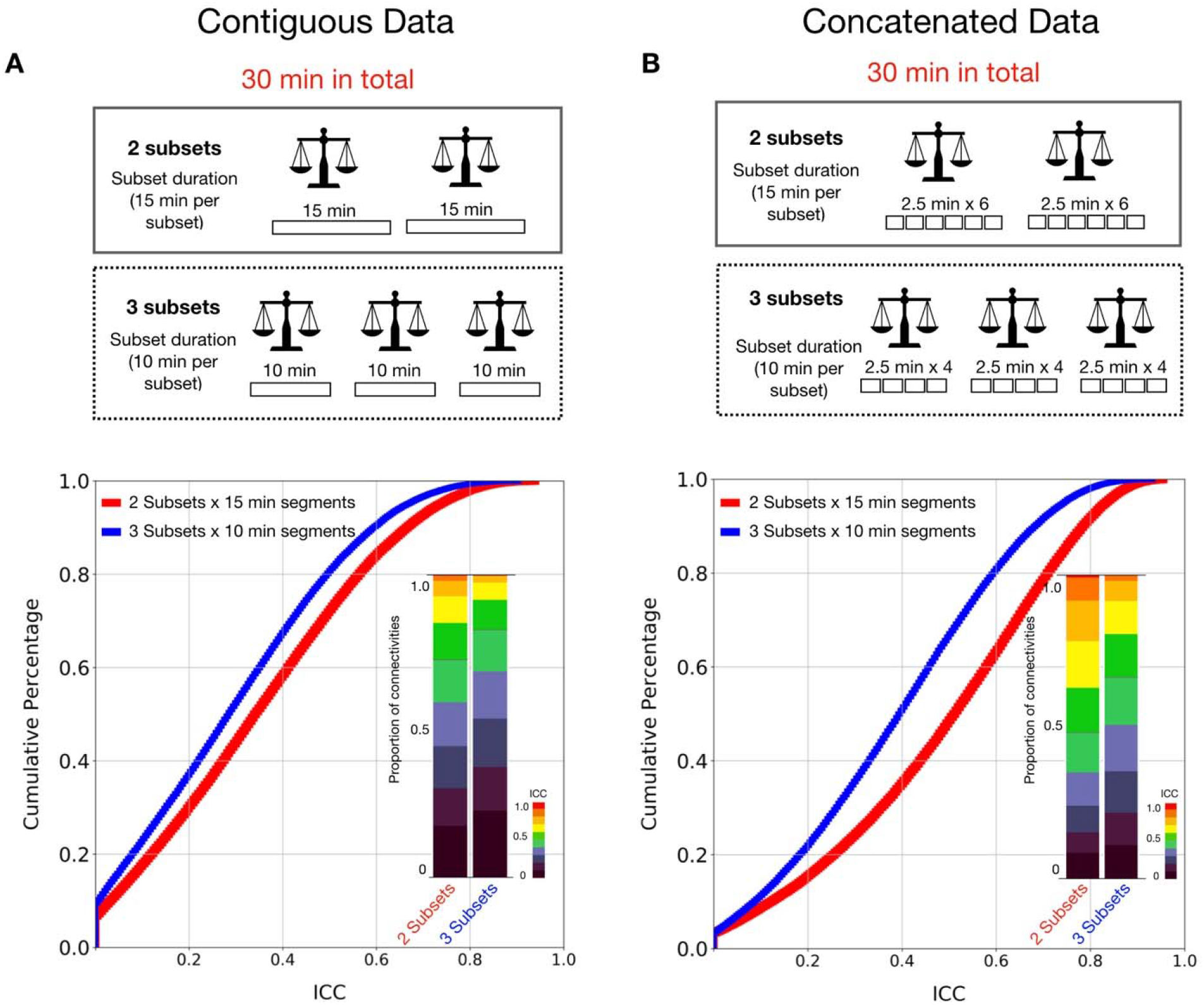
Reliability of functional connectivity estimated using less subsets yields greater reliability than using more subsets given the same total time duration. **(A)** Cumulative distribution and stacked bar plots of reliability estimates for 2 subsets x 15 min contiguous scans versus 3 subsets x 10 min scans. **(B)** Cumulative distribution and stacked bar plots of reliability estimates for 2 subsets x 15 min concatenated data versus 3 subsets x 10 min concatenated data.

We also tested subsets generated from 10, 12.5, 15, 17.5, 20, 22.5, 25, 30 min data, while the number of subsets was increased from 2 to 6 subsets (Fig S8). We observed that the reliability increased more quickly with increasing amounts of data per subset compared to increasing the number of subsets for both contiguous and concatenated data. Overall, when multiple scans were acquired, concatenating multiple scans into two subsets, instead of treating them as multiple retest subsets obtained higher ICC estimations.

## DISCUSSION

The present work evaluated the impact of different strategies for combining data from distinct fMRI scans on the reliability of functional connectivity. Several key findings emerged. First, for a given state, we found a clear advantage for combining data from multiple scans of the same state, rather than acquiring an equivalent amount of data from a single long scan. Importantly, we found that this was not necessarily true when concatenation was carried out across different fMRI states (i.e. task conditions), where we observed an advantage for concatenating segments from the same state rather than different states. When concatenation was performed across differing states, the combination from more reliable states and more homogenous states tends to yield higher reliability. We next tested whether there is an advantage for combining time-series data across scans prior to calculating functional connectivity, versus calculating the functional connectivity for each of the individual scans and then averaging; we found no clear evidence of superiority for either strategy. Finally, as a growing number of high characterization test-retest designs are emerging, we drew attention to basic design principles to be considered in the design of test-retest studies.

The present work is not the first to suggest an advantage for concatenation from multiple shorter scans over a single long scan. For example, previous work using the MyConnectome dataset has shown that functional connectivity measurements in a single subject converge faster with multiple scans of shorter length than with a single longer scan (Laumann et al., 2015). One possible reason is that the noise obtained from the scanner (e.g. pink noise or 1/f noise) might be flattened out when concatenating multiple scans together (Afshinpour et al., 2008; Akhrif et al., 2018). When a single long scan is acquired, however, the noise is maintained as the main variation between test and retest sessions, thereby reducing reliability. Another possibility is that the participants might have low head motion at the beginning of each scan (i.e., an onset effect). A recent study found that head motion events occur more frequently with longer scan length and suggested a scanner break inside the scanner to improve head motion in both children and adults samples (Meissner et al., 2020). The present work tested for the presence of an onset effect (i.e., whether concatenating time-series data from the onset of each scan could achieve higher reliability) but did not find any clear evidence to support this hypothesis. This result is consistent with previous findings that the reliability is reduced when truncating the time points from the end of the time series (Shah et al., 2016). Nevertheless, the overall results highlight the advantages of concatenation in the calculation of functional connectivity and suggest acquiring multiple shorter scans instead of a single long scan in study design--an approach adopted by the recent NIH ABCD Study (Casey et al., 2018).

It is important to note that the concatenation method itself (i.e. cutting the time-series into shorter contiguous segments and z-scoring the segments) on the same scan did not show any benefits for reliability. This suggests that the non-stationary noise, if any, might not be removed by simply Z-scoring a short contiguous time-series. In fact, our results showed that cutting the original timeseries into shorter segments *decreases* reliability (Fig. S2C). These findings indicate the downside of the concatenating shorter scans--a potential loss of the temporal continuity (e.g. auto-correlation) of the fMRI signals, which appear to be important (Arbabshirani et al., 2019; Kumar et al., 2016). One other downside of opting to concatenate multiple shorter scans, rather than pursue a single long scan, is the potential limitations this strategy introduces on temporal dynamic analyses--particularly those focused on the frequency domain (Hutchison et al., 2013). Thus, in the end, it is fair to say that there are benefits to obtaining multiple shorter scans and concatenating them, though the trade-offs should be carefully considered during experimental design.

Aside from the experimental design, our findings suggest that while concatenating scans from multiple scans to maximize data is clearly beneficial, the increases in reliability achieved will be dependent in part by the reliability of the individual scan states being combined and the homogeneity among states used in hybrid combination. Researchers usually pay attention to the total amount of data merged, yet how the data has been merged might be less heeded. In the current study, we have systematically examined differing strategies for generating hybrid data. In contrast to prior suggestions (Elliott et al., 2019), we found that when matched the number of segments in concatenation strategy, the generation of a hybrid dataset from different states does not necessarily produce functional connectivity measures with higher reliability than those obtained during resting state. In our analyses, concatenating resting state with the same concatenation strategy actually showed a substantially higher reliability than most of the hybrid combinations. This could be because merging different states introduces more variations leading to lower reliability. We have examined this hypothesis and found supportive evidence that hybrid data from fewer states are more likely to yield higher reliability than that from more different states. When concatenating multiple states data, the homogeneity between states determines the reliability of hybrid data. Overall, concatenating data from more reliable states, more homogeneous (i.e., high similarity of FC between states, same task scans, or only resting scans) showed higher reliability than data combined across states (i.e. hybrid). With the equivalent number of concatenation scans, concatenating from more reliable states trends to yield higher reliability.

Another important finding of the proposed work was the seeming similarity of results when concatenating time-series data prior to calculating the functional connectivity matrix, and the alternative of averaging functional connectivity matrices across independent scans. While the former strategy can be more memory-intensive, the latter is more computationally intensive. Depending on the length of the time-series data per scan, researchers may prefer to implement one way or the other. Our results suggest that the two methods should yield roughly equivalent functional connectivity matrices. However, concatenating the time-series first appears to have an advantage when the number of parcels was relatively small. A related question that arises when designing and analyzing highly characterized samples of individuals, is whether to use the maximum amount of data available for the ‘test’ and ‘retest’ subsets, which would be split halves of the total data, or to make the individuals scans the test unit and have multiple retest subsets. Our analyses suggest that unless data amounts are more copious than nearly all acquired to date, the aggregation of data into two halves will provide the most reliable test- and retest units (Noble et al., 2017). The alternative approach would provide the tightest confidence intervals for reliability estimation, but as shown in our analyses, the actual reliability will be undesirably low (Zou, 2012). Of note, the subcortical regions and cerebellum might require more data to achieve a similar level of reliability than cortical regions (Noble et al., 2017). In the present work, the subcortical regions and cerebellum are not yet included and will be important to characterize in future work.

In addition, the current work examined preprocessing options that need to be considered in functional connectivity analyses carried out in fMRI analysis across states - global signal regression and task regression. Our findings demonstrated that global signal regression is the main preprocessing step that dominates the functional connectivity pattern (Fig 7E). Nevertheless, the determinants of reliability for hybrid data (i.e. reliability and homogeneity of states used for hybrid combination) are replicable with or without either of these two steps; the reliability of each state and the homogeneity between states used in hybrid determines the reliability of the hybrid data. We also identified the presence of an interaction between GSR and ICA-FIX for resting-state data; the observed advantage for no GSR with resting-state data is not observed when ICA-FIX was carried out. Previous studies have demonstrated that ICA-FIX enables noise removal, though can leave substantial spatial artifacts (Burgess et al., 2016). A combination with GSR appears to clean up such artifacts and improve brain-behavior prediction (Burgess et al., 2016; Li et al., 2019). This finding echoed previous work and suggested the value of adding GSR process if ICA-FIX is performed for concatenated resting data. In addition, it is worth noting that the task and corresponding hybrid data without task regression showed higher reliability than that with task regression. Our finding that reliability was higher for resting-state fMRI data than those obtained with any of the six tasks when task-regression was used, makes the impact clear. However, an obvious consideration is that the higher FC reliability might be driven, at least in part, by the consistent task-evoked activation in individuals (Cole et al., 2019). In particular, for blocked task activation design (e.g. the HCP tasks used here), the block-wise temporal fluctuation from the task-evoked hemodynamic response might be dominant in connectivity estimates if the task-induced activation is not removed. Previous studies have shown that task-evoked coactivation might induce an inflation effect on estimates of intrinsic functional connectivity (Duff et al., 2018). As such, stimulus-evoked activity is suggested to clean up the ‘background connectivity’ in the task paradigm with a focus on intrinsic FC (Fair et al., 2007; Ito et al., 2019; Lurie et al., 2020; Norman-Haignere et al., 2012).

Of note, our study also examined the functional connectivity using partial correlation. We revealed a similar advantage for combining data from multiple scans of the same state over a single long scan using partial correlation. However, we did not apply the regularized estimation (e.g. L1-norm regularization) in the present study (Smith et al., 2013, 2011). Given that the efficient calculation of partial correlation for large-scale connectivity relies on the shrinkage estimation and appropriate regularization, it will be important for future studies to further examine the optimal regularization parameters for concatenated data.

Finally, it is important to note that reliability is not the only criterion for valid neuroimaging studies toward individual investigations and clinical neuroscience. The classification and behavioral prediction also depend on the validity of the measurement, for example, the extent to which measurement assesses the desired targets (i.e. construct validity) and whether the measurement content covers the domain to be measured (i.e. content validity). A recent study has demonstrated that the task activation improves the prediction but only if the in-scanner task is related to the out-scanner predicted phenotype (Greene et al., 2020). Thus, depending on the desired content, the optimal strategy (e.g. preprocessing options) and/or states (e.g. task, rest) might be different (Noble et al., 2019). In some scenarios, the denoising procedure might remove reliable artifacts and even reduce the reliability (Noble et al., 2019; Shirer et al., 2015). Nevertheless, when no specific content of the phenotype to be targeted, optimizing the reliability is the prerequisite toward the valid measurement as reliability is the upper limit to validity (Bridgeford et al., 2020; Carmines and Zeller, 1979; Hong et al., 2020; Zuo et al., 2019). Beyond the reliability, understanding the potential impacts of data preprocessing and its nuance in determining the reliability and brain-behavior prediction needs to be carefully examined in future studies.

### Summary of Recommendations

Consistent with prior work, the findings of the present work reaffirmed the “golden rule”: <u>more data is better - whether obtained in a single condition or scan session, or multiple</u>. With that said, our findings allowed us to make a base set of recommendations for consideration in functional connectivity study design and data analysis, when the amount of data available is equal. Specifically, we suggest that when possible, researchers:

1. Acquire and concatenate multiple short scans or sessions, rather than a single long scan. *Caveat:* This recommendation is appropriate for studies focused on static functional connectivity, not dynamic. For the latter, the suitability of multiple scans for analysis is dependent on the specific algorithms employed.
2. Concatenation of multiple scans with shorter duration offers higher reliability than those with a longer duration. Thus, a shorter duration between scans is recommended for multiple scans design. *Of note*, data need to be acquired from multiple scans; back-to-back shorter segments that are split from a single long scan does not appear to have any advantage.
3. Concatenation of data within a single scan condition is more likely to offer higher reliability than those across multiple conditions. Among conditions, concatenating more reliable conditions, and more homogeneous conditions (i.e. high similarity of functional connectivity between conditions) is more likely to yield higher reliability.
4. When conducting test-retest reliability studies, after multiple scans are acquired, the scans should be concatenated into two subsets, instead of treated as multiple.
5. When comparing the reliability, beyond the total amount of data, be cautious with the strategies of how data is concatenated for a fair comparison.

## Conclusion

This work systematically evaluated the impact of data concatenation strategies on test-retest reliability of functional connectivity. When looking at equivalent amounts of data, our results highlight the potential advantages of concatenated data relative to contiguous, and other factors that can lead to higher or lower reliability for concatenated data. We drew attention to dependencies of the resultant reliability of a concatenated scan on the individual scans from which data is aggregated. Finally, we drew attention to the differential impact of preprocessing (i.e. task regression, global signal regression). Combined, the present work provides an overview of multiple decisions that should be considered in efforts to optimize reliability during the acquisition and analysis of functional connectivity data.

## ACKNOWLEDGEMENT

This work was supported by gifts from Joseph P. Healey, Phyllis Green, and Randolph Cowen to the Child Mind Institute and the National Institutes of Health (Brain Research through Advancing Innovative Neurotechnologies Initiative Grant Nos. R01-MH111439 [to MPM and Charles E. Schroeder], R24MH114806 [to MPM], and R01MH120482 [to MPM]). Additional grant support for JTV comes from 1R01MH120482-01 (to Theodore D. Satterthwaite, MPM) and funding from Microsoft Research. We thank Russell Takeshi Shinohara and Seok-Jun Hong for valuable discussions and suggestions.

