## Supplemental figures and tables for "Impact of Concatenating fMRI Data on Reliability for Functional Connectomics"

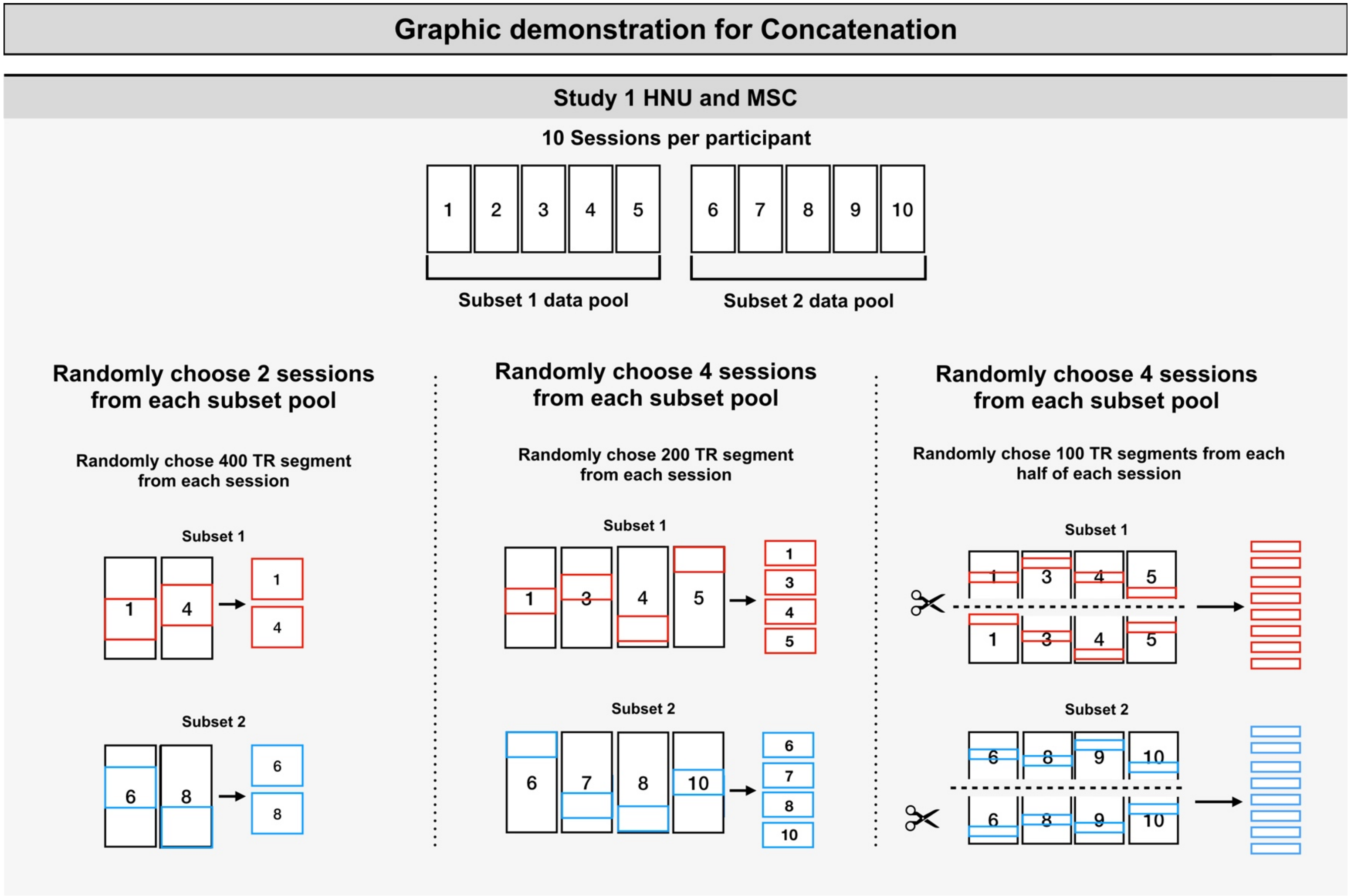


**Figure S1. Graphic detailing segment selection and concatenation strategies used for HNU and MSC datasets for Study-1.**

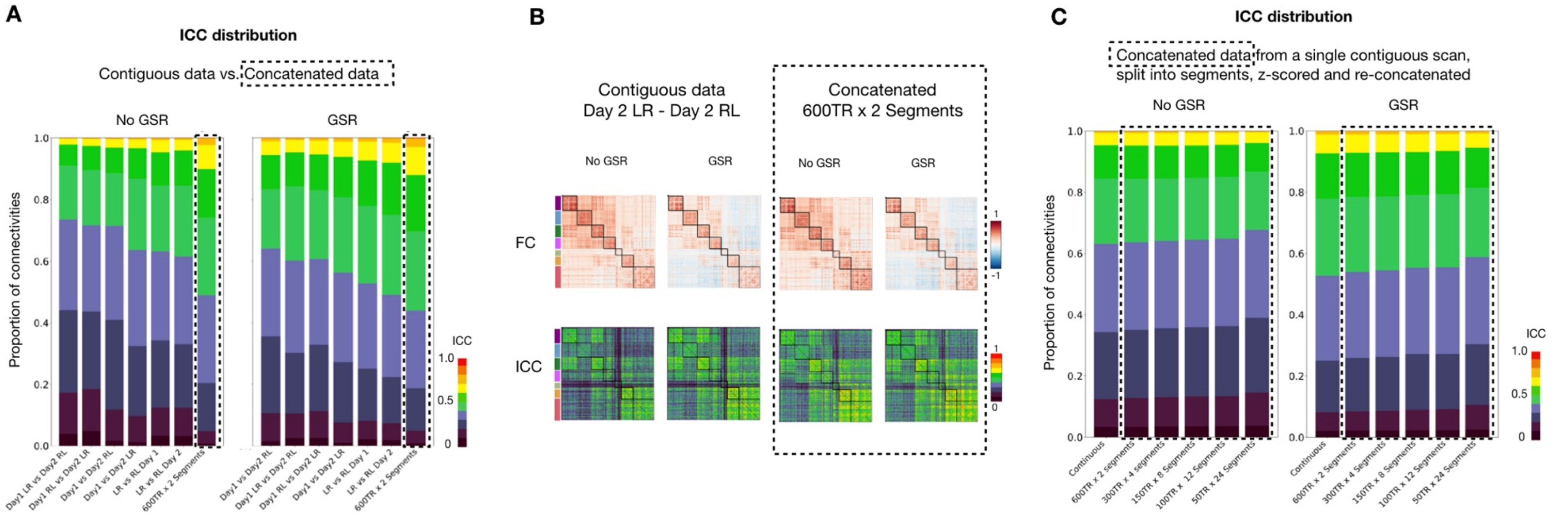


**Figure S2. Reliability of edgewise functional connectivity of data concatenated from multiple different scans and the same scan.** **(A)** The ICC distribution of functional connectivity was plotted as stacked bar plots in HCP dataset with and without GSR preprocessing step. **(B)** The average functional connectivity matrix and ICC matrix for contiguous resting-state (top) and concatenated resting-state data (bottom). **(C)** Stacked bar plots of ICC distribution for functional connectivity in contiguous time-series (1200 time points) and when the same time-series was cut into segments, z-scored, and re-concatenated together.

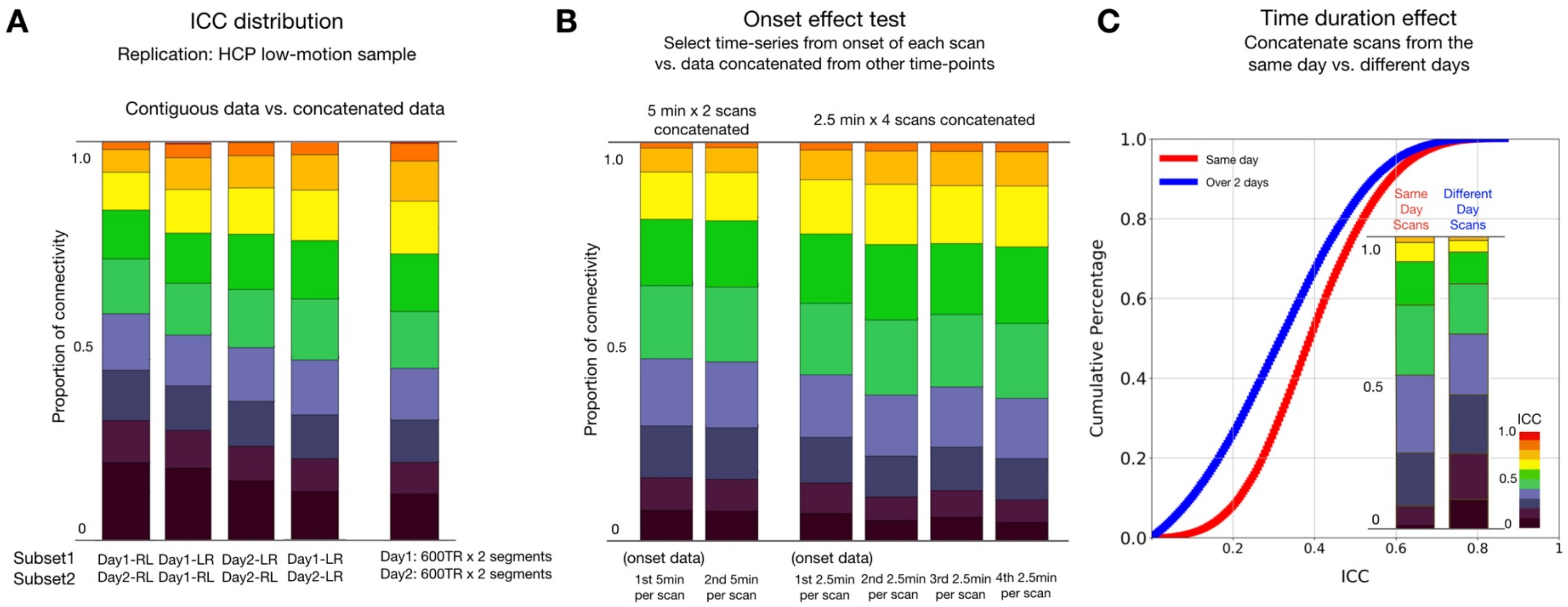


**Figure S3. The distribution of reliability for functional connectivity. (A) The concatenation effects in a low-motion HCP sample. (B)** Concatenating time-series data from the onset of each scan have no clear advantages to achieve higher reliability than concatenated data from other time-points. **(C)** Effect of concatenating segments from the same day (same session) compared to concatenating segments from two different days. Reliability of FC calculated from same-day segments were greater than that of FC calculated over 2 days.

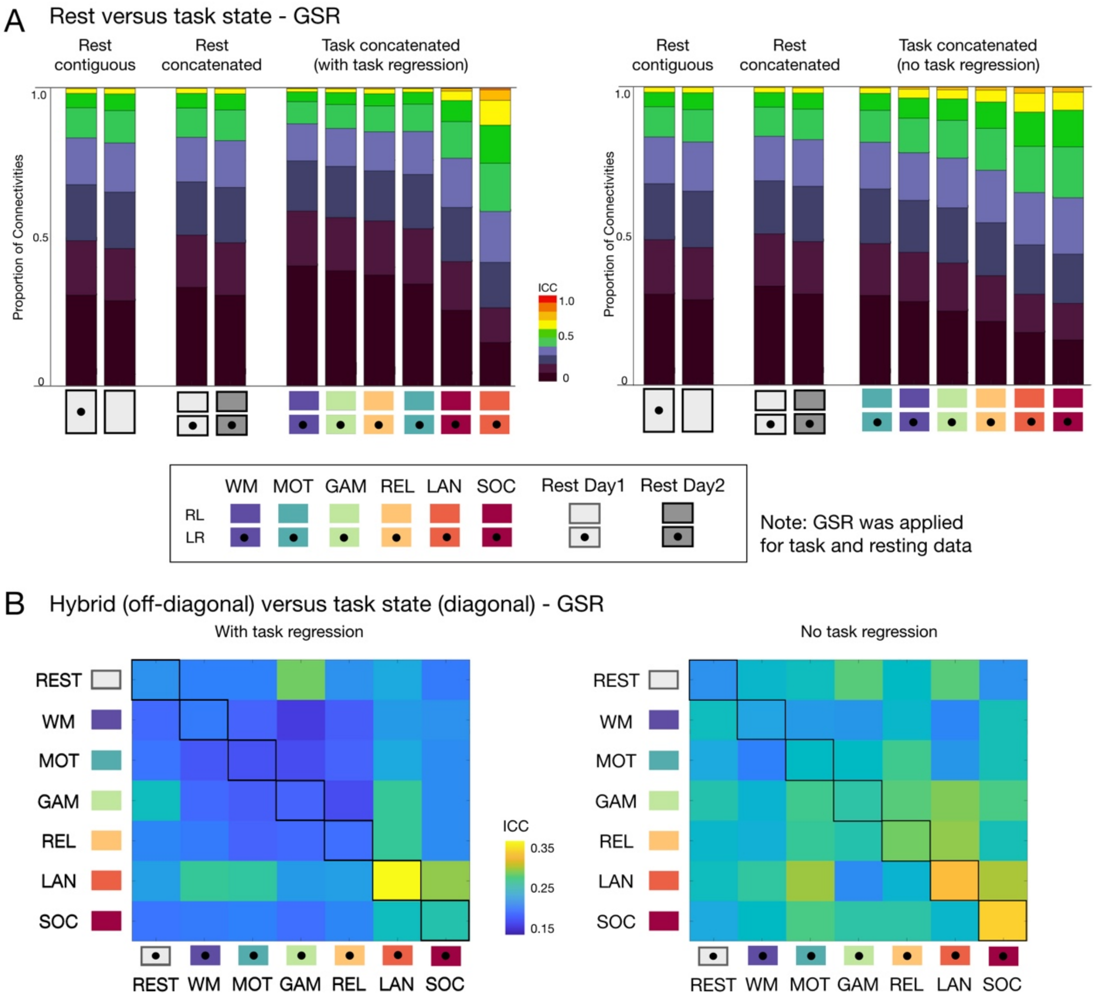


**Figure S4. Reliability of edgewise functional connectivity measures calculated using each, resting-state, task and hybrid fMRI data with GSR.** **(A)** Proportion of connections with poor, moderate, good and excellent ICC when calculated for resting state and each of six task conditions in the HCP dataset, with (left-panel) and without task regression (right-panel). **(B)** Mean ICC for functional connectivity measures based on hybrid data generated by concatenating two scans from two different task states (off-diagonal) and the same task state (diagonal). See results without GSR in Fig 2.

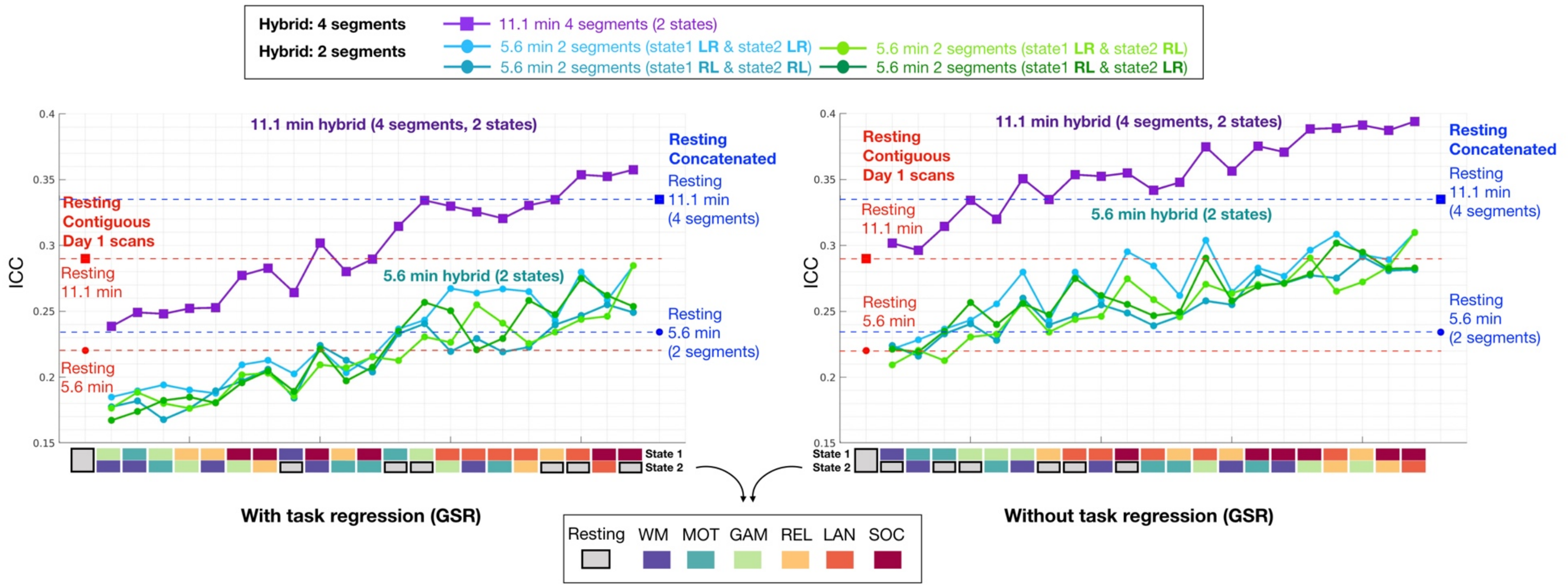


**Figure S5. Reliability of edgewise functional connectivity measures calculated for resting and hybrid data generated from 2 tasks using either 2 or 4 segments with GSR.** The same length of contiguous resting data was truncated from LR and RL scans of the Day 1 session (red dots, 5.6 min; red square: 11.1 min). The concatenated resting data was generated from 2 scans (LR and RL scans from Day 1) and 4 scans (LR, RL scans from Day 1 and Day 2). See results without GSR in Fig 4.

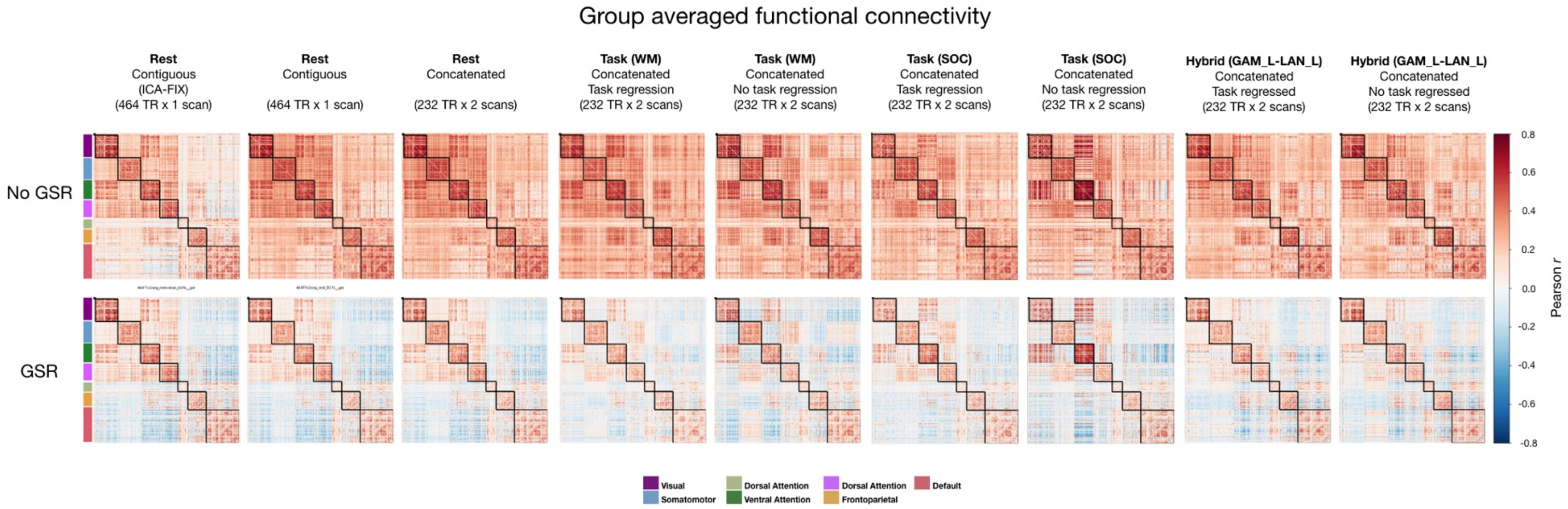


**Figure S6. Group averaged functional connectivity matrices with different preprocessing options.** Functional connectivity matrices shown for data processed without GSR (top row) and with GSR (bottom row).

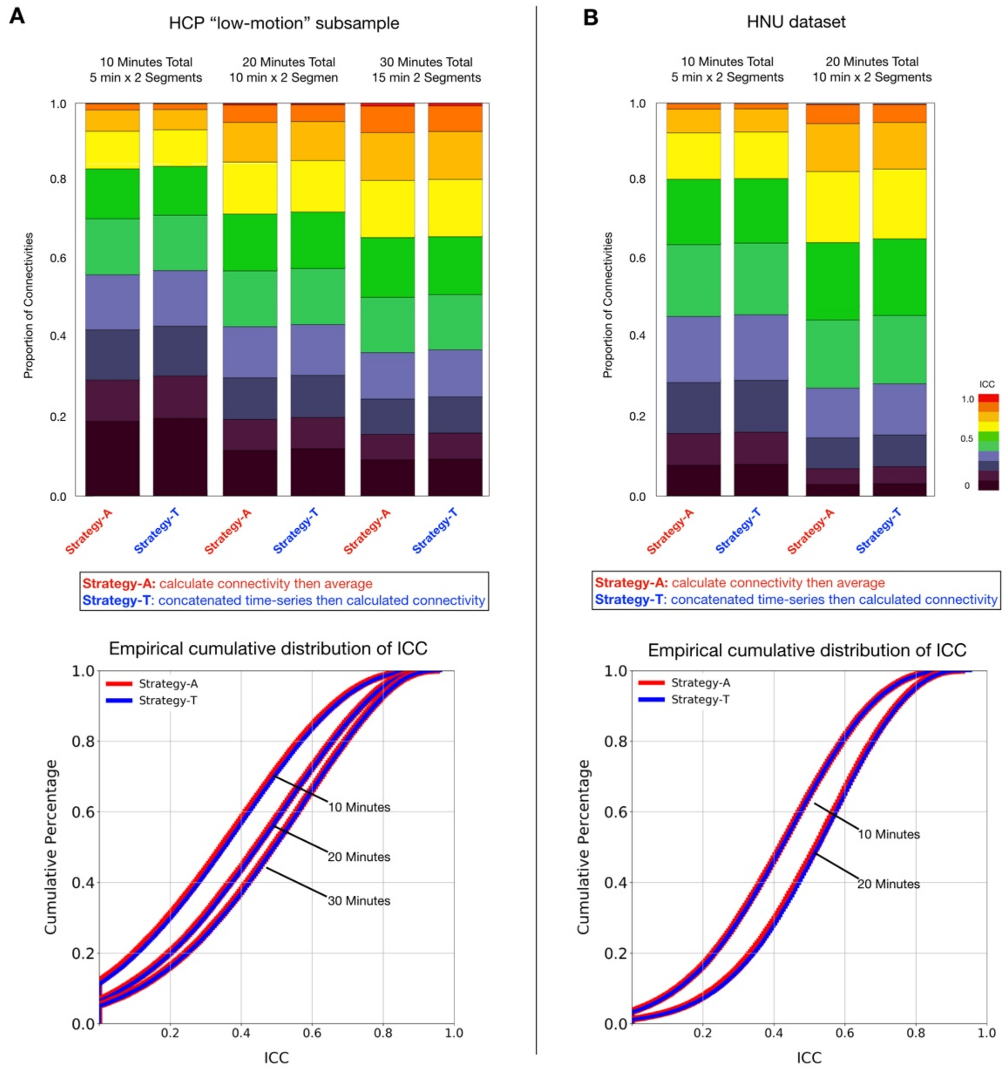


**Figure S7. Calculation of edgewise functional connectivity measures for scans individually and then averaging, versus calculation after concatenation of scans.** The strategy calculating the FC of each segment first then average the FC as the final FC for each subset was denoted as Strategy-A, while concatenating time-series of all segments then calculating FC per subset as Strategy-T. The bar stacked plots of ICC distribution were showed in HCP low-motion subsample (A) and HNU dataset (B).

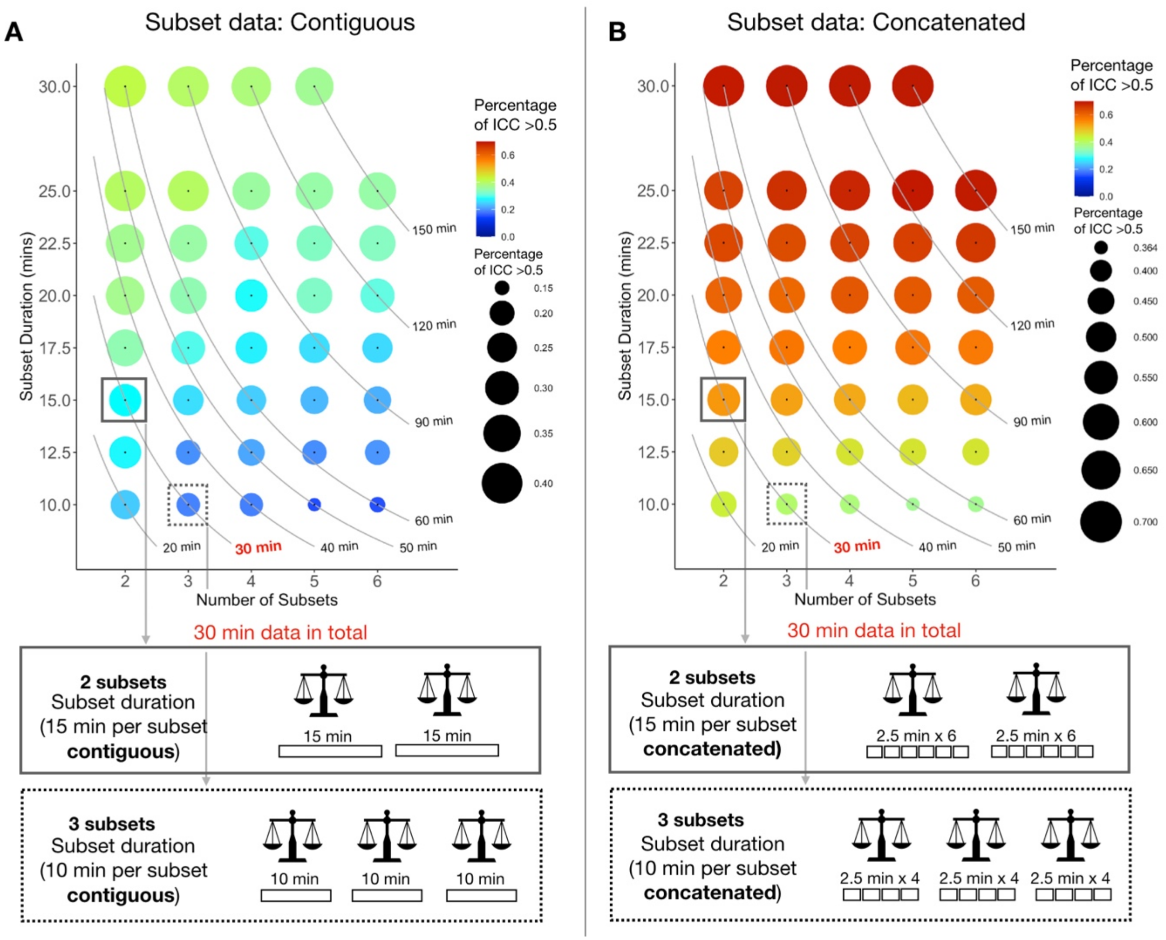


**Figure S8. Reliability of functional connectivity estimated across different numbers of subsets and different length of data within each subset.** Bubbles represent the ratio of ICC values > 0.5 at each combination of the number of test-retest subsets (x-axis) by the duration for each subset (y-axis). The contour line of total amount of time (subsets x duration) are plotted in gray lines. **(A)** Each subset data was created from a contiguous scan while each subset in **(B)** was created from concatenation of shorter segments from multiple scans. Two example combinations are marked by rectangles (solid line: 2 subsets with 15 min data per subset, dashed line: 3 subsets with 10 min data per subset).

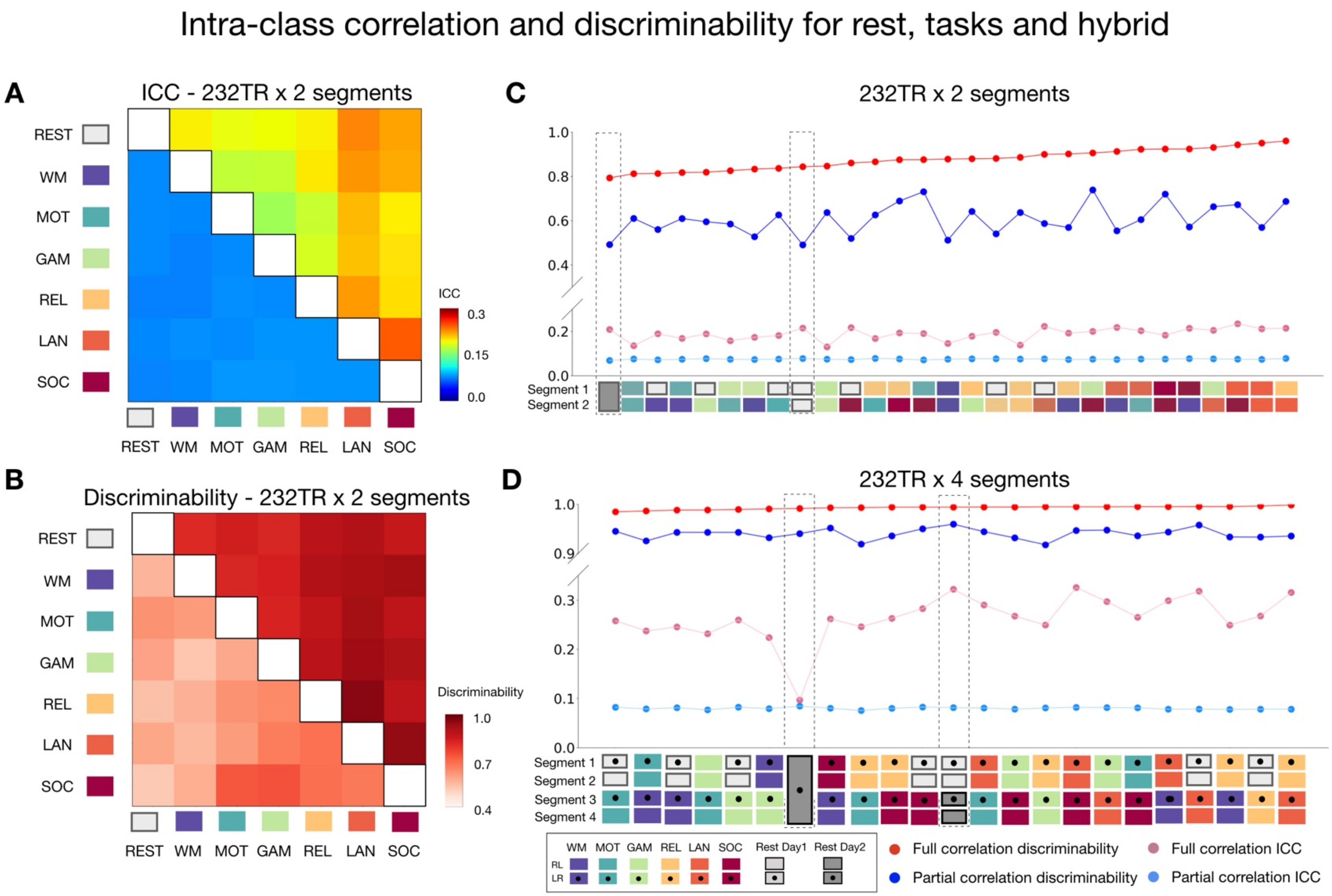


**Figure S9. Comparison of intra-class correlation (ICC) and discriminability of functional connectivity calculated from full correlation and partial correlation based on Schaefer100 parcellation.** Upper triangles show results for ICC **(A)** and discriminability **(B)** of hybrid FC calculated using Pearson correlation. Lower triangles show ICC and discriminability of hybrid FC calculated from partial correlation. Discriminability and ICC of FC calculated using Pearson correlation (red shades) and partial correlation (blue shades) for rest, tasks and hybrid data concatenated with 2 segments **(C)** and 4 segments **(D).** Dash line boxes highlighted the contiguous and concatenated resting condition.

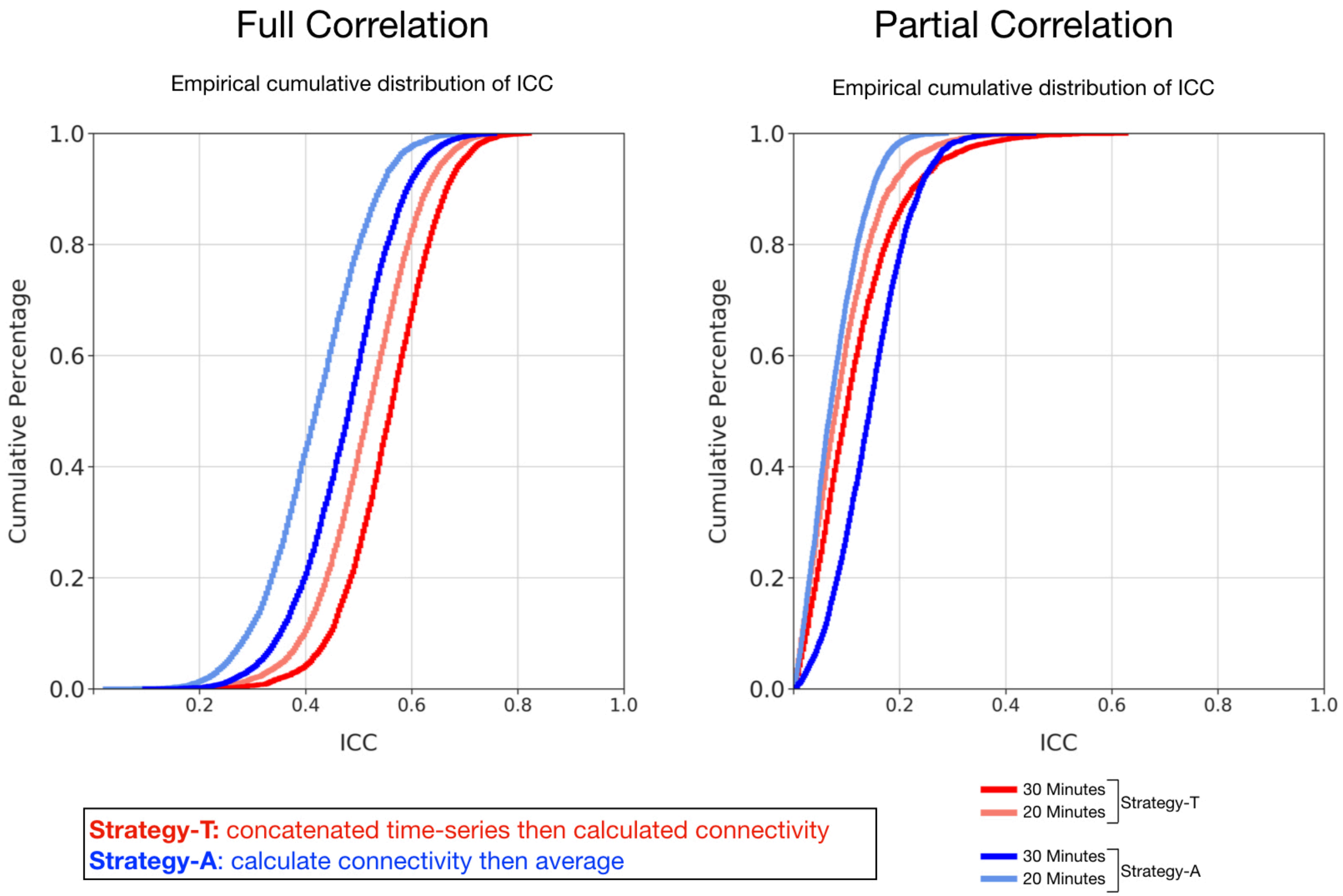


**Figure S10. Comparison of Strategy-T and Strategy-A for FC created using partial correlations based on Schaefer100 parcellation.** Reliability of edges from FC calculated with partial correlation were higher for Strategy-A (average multiple FC before reliability calculation) than with Strategy-T (concatenate first then average) but opposite for Pearson correlation.

Table S1. The mean and standard deviation of the head motion (Framewise displacement) for dataset used in the current study.

| Task | HCP | HCP Retest Pre | HCP Retest Post | HCP 22 Subject Subsample |
|---|---|---|---|---|
| REST1 | 0.134 (0.027) | 0.121 (0.017) | 0.121 (0.017) | 0.121 (0.017) |
| REST2 | 0.137 (0.026) | 0.122 (0.016) | 0.122 (0.016) | 0.122 (0.016) |
| WM | 0.137 (0.032) | 0.121 (0.019) | 0.121 (0.019) | 0.121 (0.019) |
| GAM | 0.133 (0.029) | 0.119 (0.018) | 0.119 (0.018) | 0.119 (0.018) |
| LAN | 0.143 (0.041) | 0.123 (0.018) | 0.123 (0.018) | 0.123 (0.018) |
| MOT | 0.148 (0.034) | 0.123 (0.017) | 0.123 (0.017) | 0.123 (0.017) |
| SOC | 0.139 (0.038) | 0.119 (0.019) | 0.119 (0.019) | 0.119 (0.019) |
| REL | 0.143 (0.033) | 0.123 (0.018) | 0.123 (0.018) | 0.123 (0.018) |
| **Task** | **MSC** | **HNU** | | |
| Rest | 0.134 (0.027) | 0.121 (0.017) | | |

HCP “low-motion” subsample IDs
100307 106521 111413 111716 118730 122620 127933 129028 136227 147737 148335
151627 159239 162228 173940 181232 201818 203418 239944 552544 679568 782561

Table S2. The averaged test-retest reliability of the edgewise connections for task and resting state preprocessed with different preprocessing options (i.e. GSR, task-regression).

| | No GSR | | GSR | |
|---|---|---|---|---|
| | 2 segments, 5.6 min in total | | | |
| Task | With task regression | Without task regression | With task regression | Without task regression |
| GAM | 0.185 (0.164) | 0.270 (0.189) | 0.188 (0.171) | 0.286 (0.196) |
| WM | 0.186 (0.163) | 0.243 (0.186) | 0.179 (0.166) | 0.250 (0.19) |
| REL | 0.219 (0.175) | 0.305 (0.2) | 0.197 (0.175) | 0.310 (0.201) |
| MOT | 0.217 (0.169) | 0.262 (0.178) | 0.206 (0.17) | 0.245 (0.182) |
| SOC | 0.256 (0.182) | 0.344 (0.194) | 0.255 (0.187) | 0.357 (0.203) |
| LAN | 0.338 (0.199) | 0.338 (0.199) | 0.348 (0.207) | 0.348 (0.207) |

| Rest Conditions | No GSR | GSR |
|---|---|---|
| Rest Continuous LR (Day1) | 0.219 (0.166) | 0.220 (0.174) |
| Rest Continuous RL (Day1) | 0.230 (0.174) | 0.228 (0.177) |
| Rest Concatenated Day 1 | 0.226 (0.175) | 0.236 (0.184) |
| Rest Concatenated Day 2 | 0.228 (0.171) | 0.228 (0.178) |

Table S3. The averaged test-retest reliability of the edgewise connections for resting state and hybrid combinations with two different tasks preprocessed with different preprocessing options (i.e. GSR, task-regression, and ICA-FIX).

| No GSR | | With task regression | | | | | Without task regression | | | | |
|---|---|---|---|---|---|---|---|---|---|---|---|
| | | 2 segments, 5.6 min in total | | | | 4 segments, 11.1 min in total | 2 segments, 5.6 min in total | | | | 4 segments, 11.1 min in total |
| Task1 | Task2 | 1L, 2L | 1R, 2R | 1L, 2R | 1R, 2L | 1L, 1R, 2L, 2R | 1L, 2L | 1R, 2R | 1L, 2R | 1R, 2L | 1L, 1R, 2L, 2R |
| GAM | LAN | 0.270 (0.185) | 0.234 (0.177) | 0.231 (0.179) | 0.239 (0.182) | 0.326 (0.199) | 0.299 (0.189) | 0.264 (0.182) | 0.263 (0.183) | 0.274 (0.187) | 0.361 (0.200) |
| GAM | MOT | 0.208 (0.169) | 0.184 (0.160) | 0.194 (0.164) | 0.185 (0.163) | 0.268 (0.184) | 0.257 (0.181) | 0.238 (0.178) | 0.242 (0.176) | 0.235 (0.179) | 0.323 (0.195) |
| GAM | SOC | 0.227 (0.174) | 0.197 (0.167) | 0.208 (0.170) | 0.196 (0.168) | 0.278 (0.189) | 0.293 (0.183) | 0.271 (0.186) | 0.272 (0.185) | 0.271 (0.188) | 0.370 (0.203) |
| GAM | WM | 0.200 (0.168) | 0.192 (0.163) | 0.199 (0.166) | 0.173 (0.161) | 0.257 (0.185) | 0.270 (0.187) | 0.258 (0.185) | 0.250 (0.181) | 0.247 (0.188) | 0.335 (0.201) |
| LAN | MOT | 0.272 (0.184) | 0.234 (0.174) | 0.247 (0.180) | 0.230 (0.175) | 0.332 (0.195) | 0.285 (0.186) | 0.257 (0.178) | 0.268 (0.184) | 0.249 (0.180) | 0.355 (0.196) |
| LAN | SOC | 0.283 (0.189) | 0.255 (0.177) | 0.285 (0.184) | 0.256 (0.183) | 0.355 (0.196) | 0.304 (0.189) | 0.275 (0.180) | 0.301 (0.187) | 0.286 (0.184) | 0.384 (0.196) |
| LAN | WM | 0.264 (0.186) | 0.248 (0.177) | 0.255 (0.182) | 0.228 (0.179) | 0.331 (0.197) | 0.283 (0.195) | 0.261 (0.181) | 0.274 (0.189) | 0.248 (0.187) | 0.353 (0.203) |
| MOT | SOC | 0.234 (0.176) | 0.208 (0.167) | 0.209 (0.169) | 0.214 (0.169) | 0.296 (0.189) | 0.282 (0.179) | 0.254 (0.177) | 0.255 (0.182) | 0.269 (0.179) | 0.364 (0.195) |
| MOT | WM | 0.191 (0.163) | 0.201 (0.164) | 0.199 (0.164) | 0.183 (0.161) | 0.268 (0.181) | 0.225 (0.177) | 0.243 (0.177) | 0.224 (0.173) | 0.224 (0.177) | 0.311 (0.193) |
| REL | GAM | 0.212 (0.171) | 0.189 (0.162) | 0.186 (0.165) | 0.190 (0.165) | 0.274 (0.186) | 0.286 (0.189) | 0.291 (0.194) | 0.261 (0.186) | 0.272 (0.19) | 0.375 (0.206) |
| REL | LAN | 0.272 (0.186) | 0.230 (0.175) | 0.233 (0.177) | 0.247 (0.182) | 0.336 (0.197) | 0.297 (0.193) | 0.267 (0.185) | 0.263 (0.183) | 0.282 (0.189) | 0.372 (0.201) |
| REL | MOT | 0.215 (0.170) | 0.199 (0.164) | 0.200 (0.167) | 0.193 (0.165) | 0.292 (0.188) | 0.265 (0.184) | 0.254 (0.181) | 0.255 (0.182) | 0.245 (0.180) | 0.357 (0.199) |
| REL | SOC | 0.226 (0.176) | 0.208 (0.169) | 0.203 (0.170) | 0.201 (0.168) | 0.286 (0.190) | 0.293 (0.186) | 0.273 (0.184) | 0.276 (0.189) | 0.265 (0.186) | 0.377 (0.201) |
| REL | WM | 0.205 (0.170) | 0.200 (0.167) | 0.197 (0.169) | 0.182 (0.163) | 0.277 (0.188) | 0.270 (0.193) | 0.274 (0.186) | 0.263 (0.190) | 0.262 (0.192) | 0.365 (0.206) |
| SOC | WM | 0.214 (0.173) | 0.231 (0.175) | 0.218 (0.170) | 0.205 (0.170) | 0.295 (0.191) | 0.276 (0.185) | 0.272 (0.184) | 0.274 (0.181) | 0.260 (0.186) | 0.367 (0.197) |

| | ICAFIX | | | | | No ICAFIX | | | | |
|---|---|---|---|---|---|---|---|---|---|---|
| Rest Conditions | LR | | RL | | 1L, 1R, 2L, 2R | LR | | RL | | 1L, 1R, 2L, 2R |
| Rest Contiguous (Day1), 5.6min | 0.204 (0.172) | 0.227 (0.177) | 0.204 (0.172) | | -- | 0.219 (0.166) | 0.230 (0.174) | 0.219 (0.166) | | -- |
| Rest Concatenated (Day1, Day2), 5.6min | 0.239 (0.180) | 0.238 (0.180) | 0.239 (0.180) | | -- | 0.243 (0.173) | 0.233 (0.174) | 0.243 (0.173) | | -- |
| Rest Contiguous (Day1), 11.1min | 0.247 (0.184) | 0.270 (0.193) | 0.247 (0.184) | | -- | 0.226 (0.174) | 0.296 (0.188) | 0.226 (0.174) | | -- |
| Rest Concatenated - 4 scans, 11.1min | -- | | -- | | 0.343 (0.199) | -- | | -- | | 0.338 (0.188) |

Table S3 continued

| GSR | | With task regression | | | | | Without task regression | | | | |
|---|---|---|---|---|---|---|---|---|---|---|---|
| | | 2 segments, 5.6 min in total | | | | 4 segments, 11.1 min in total | 2 segments, 5.6 min in total | | | | 4 segments, 11.1 min in total |
| Task1 | Task2 | 1L, 2L | 1R, 2R | 1L, 2R | 1R, 2L | 1L, 1R, 2L, 2R | 1L, 2L | 1R, 2R | 1L, 2R | 1R, 2L | 1L, 1R, 2L, 2R |
| GAM | WM | 0.185 (0.168) | 0.177 (0.163) | 0.176 (0.166) | 0.167 (0.162) | 0.238 (0.190) | 0.279 (0.194) | 0.260 (0.191) | 0.256 (0.190) | 0.256 (0.195) | 0.350 (0.21) |
| GAM | MOT | 0.194 (0.170) | 0.167 (0.159) | 0.180 (0.166) | 0.182 (0.166) | 0.248 (0.191) | 0.255 (0.186) | 0.228 (0.181) | 0.232 (0.180) | 0.240 (0.185) | 0.320 (0.203) |
| MOT | WM | 0.189 (0.168) | 0.182 (0.164) | 0.188 (0.165) | 0.174 (0.161) | 0.249 (0.188) | 0.228 (0.185) | 0.216 (0.175) | 0.220 (0.176) | 0.219 (0.180) | 0.296 (0.201) |
| REL | WM | 0.187 (0.170) | 0.189 (0.169) | 0.180 (0.167) | 0.180 (0.167) | 0.252 (0.195) | 0.283 (0.197) | 0.279 (0.190) | 0.270 (0.195) | 0.269 (0.194) | 0.375 (0.211) |
| REL | GAM | 0.190 (0.171) | 0.176 (0.164) | 0.176 (0.164) | 0.185 (0.169) | 0.252 (0.193) | 0.293 (0.194) | 0.291 (0.198) | 0.272 (0.191) | 0.295 (0.195) | 0.391 (0.209) |
| REL | MOT | 0.202 (0.173) | 0.184 (0.164) | 0.185 (0.165) | 0.189 (0.167) | 0.264 (0.192) | 0.262 (0.185) | 0.246 (0.180) | 0.245 (0.180) | 0.249 (0.185) | 0.347 (0.203) |
| GAM | SOC | 0.209 (0.177) | 0.197 (0.17) | 0.201 (0.173) | 0.195 (0.173) | 0.277 (0.198) | 0.296 (0.188) | 0.277 (0.196) | 0.290 (0.192) | 0.278 (0.193) | 0.388 (0.207) |
| SOC | WM | 0.203 (0.173) | 0.213 (0.173) | 0.207 (0.175) | 0.197 (0.169) | 0.280 (0.196) | 0.276 (0.188) | 0.271 (0.187) | 0.271 (0.186) | 0.271 (0.192) | 0.370 (0.205) |
| REL | SOC | 0.212 (0.176) | 0.206 (0.172) | 0.203 (0.171) | 0.204 (0.174) | 0.282 (0.198) | 0.289 (0.188) | 0.281 (0.190) | 0.283 (0.191) | 0.282 (0.187) | 0.387 (0.205) |
| MOT | SOC | 0.215 (0.177) | 0.204 (0.169) | 0.215 (0.173) | 0.207 (0.172) | 0.289 (0.196) | 0.264 (0.182) | 0.255 (0.182) | 0.263 (0.187) | 0.258 (0.180) | 0.356 (0.201) |
| LAN | MOT | 0.267 (0.191) | 0.219 (0.176) | 0.241 (0.185) | 0.229 (0.181) | 0.320 (0.205) | 0.284 (0.194) | 0.239 (0.18) | 0.259 (0.187) | 0.246 (0.186) | 0.342 (0.207) |
| LAN | WM | 0.263 (0.191) | 0.229 (0.180) | 0.255 (0.189) | 0.220 (0.181) | 0.325 (0.206) | 0.295 (0.202) | 0.248 (0.187) | 0.274 (0.196) | 0.255 (0.193) | 0.355 (0.213) |
| GAM | LAN | 0.267 (0.194) | 0.219 (0.179) | 0.226 (0.185) | 0.250 (0.188) | 0.329 (0.209) | 0.304 (0.195) | 0.258 (0.190) | 0.270 (0.191) | 0.290 (0.195) | 0.374 (0.208) |
| REL | LAN | 0.265 (0.189) | 0.222 (0.181) | 0.225 (0.181) | 0.258 (0.189) | 0.330 (0.206) | 0.308 (0.198) | 0.275 (0.191) | 0.265 (0.189) | 0.301 (0.195) | 0.389 (0.210) |
| LAN | SOC | 0.284 (0.192) | 0.249 (0.182) | 0.284 (0.188) | 0.253 (0.187) | 0.357 (0.204) | 0.31 (0.192) | 0.281 (0.190) | 0.309 (0.193) | 0.283 (0.189) | 0.394 (0.205) |
| | | ICAFIX | | | | | No ICAFIX | | | | |
| Rest Conditions | | LR | | RL | | 1L, 1R, 2L, 2R | LR | | RL | | 1L, 1R, 2L, 2R |
| Rest Contiguous (Day1), 5.6min | | 0.227 (0.178) | 0.238 (0.185) | 0.227 (0.178) | | -- | 0.220 (0.175) | 0.229 (0.177) | 0.220 (0.175) | 0.229 (0.177) | -- |
| Rest Concatenated (Day1, Day2), 5.6min | | 0.248 (0.186) | 0.266 (0.190) | 0.248 (0.186) | | -- | 0.234 (0.182) | 0.239 (0.183) | 0.234 (0.182) | 0.239 (0.183) | -- |
| Rest Contiguous (Day1), 11.1min | | 0.283 (0.192) | 0.297 (0.203) | 0.283 (0.192) | | -- | 0.290 (0.187) | 0.279 (0.189) | 0.290 (0.187) | 0.279 (0.189) | -- |
| Rest Concatenated - 4 scans, 11.1min | | -- | | -- | | 0.360 (0.207) | -- | | -- | | 0.335 (0.203) |

Table S4. The averaged test-retest reliability of the edgewise connections for resting state and hybrid combinations with four different tasks preprocessed with different preprocessing options (i.e. GSR, task-regression, and ICA-FIX).

| No GSR | | | | No GSR - with task regression | | | | | | No GSR - without task regression | | | | | |
|---|---|---|---|---|---|---|---|---|---|---|---|---|---|---|---|
| Task1 | Task2 | Task3 | Task4 | 1L-2R-3L-4R | 1L-2L-3R-4R | 1L-2R-3R-4L | 1R-2R-3L-4L | 1R-2L-3L-4R | 1R-2L-3R-4L | 1L-2R-3L-4R | 1L-2L-3R-4R | 1L-2R-3R-4L | 1R-2R-3L-4L | 1R-2L-3L-4R | 1R-2L-3R-4L |
| REL | GAM | MOT | SOC | 0.259 (0.187) | 0.269 (0.187) | 0.262 (0.184) | 0.263 (0.186) | 0.267 (0.186) | 0.271 (0.184) | 0.338 (0.199) | 0.344 (0.196) | 0.335 (0.196) | 0.339 (0.200) | 0.338 (0.198) | 0.337 (0.197) |
| REL | GAM | SOC | WM | 0.264 (0.185) | 0.276 (0.189) | 0.256 (0.187) | 0.254 (0.186) | 0.273 (0.186) | 0.268 (0.186) | 0.347 (0.200) | 0.353 (0.202) | 0.346 (0.204) | 0.347 (0.207) | 0.350 (0.199) | 0.351 (0.202) |
| REL | GAM | MOT | WM | 0.265 (0.185) | 0.273 (0.186) | 0.257 (0.184) | 0.252 (0.182) | 0.269 (0.185) | 0.265 (0.186) | 0.330 (0.198) | 0.340 (0.197) | 0.330 (0.200) | 0.334 (0.202) | 0.334 (0.197) | 0.338 (0.202) |
| REL | MOT | SOC | WM | 0.274 (0.186) | 0.274 (0.188) | 0.265 (0.187) | 0.263 (0.186) | 0.273 (0.186) | 0.261 (0.185) | 0.346 (0.195) | 0.339 (0.200) | 0.336 (0.199) | 0.330 (0.199) | 0.334 (0.193) | 0.324 (0.197) |
| GAM | MOT | SOC | WM | 0.277 (0.183) | 0.278 (0.187) | 0.268 (0.187) | 0.253 (0.184) | 0.269 (0.185) | 0.255 (0.185) | 0.336 (0.192) | 0.331 (0.196) | 0.331 (0.197) | 0.324 (0.198) | 0.330 (0.195) | 0.323 (0.201) |
| REL | LAN | MOT | SOC | 0.284 (0.189) | 0.297 (0.190) | 0.289 (0.189) | 0.285 (0.190) | 0.300 (0.190) | 0.287 (0.19) | 0.332 (0.194) | 0.343 (0.194) | 0.338 (0.192) | 0.326 (0.194) | 0.338 (0.194) | 0.326 (0.197) |
| REL | GAM | LAN | MOT | 0.283 (0.189) | 0.285 (0.189) | 0.280 (0.19) | 0.285 (0.190) | 0.290 (0.189) | 0.278 (0.188) | 0.332 (0.196) | 0.338 (0.194) | 0.328 (0.193) | 0.344 (0.198) | 0.341 (0.199) | 0.332 (0.197) |
| GAM | LAN | MOT | WM | 0.286 (0.187) | 0.291 (0.188) | 0.277 (0.190) | 0.269 (0.188) | 0.287 (0.190) | 0.274 (0.190) | 0.320 (0.191) | 0.331 (0.195) | 0.323 (0.198) | 0.312 (0.196) | 0.326 (0.196) | 0.322 (0.199) |
| REL | LAN | MOT | WM | 0.287 (0.188) | 0.291 (0.190) | 0.283 (0.189) | 0.272 (0.187) | 0.290 (0.190) | 0.285 (0.192) | 0.328 (0.193) | 0.339 (0.198) | 0.330 (0.198) | 0.319 (0.197) | 0.335 (0.194) | 0.334 (0.201) |
| GAM | LAN | MOT | SOC | 0.285 (0.191) | 0.299 (0.19) | 0.29 (0.188) | 0.285 (0.191) | 0.294 (0.192) | 0.279 (0.189) | 0.329 (0.196) | 0.339 (0.195) | 0.335 (0.192) | 0.330 (0.194) | 0.338 (0.198) | 0.325 (0.197) |
| REL | GAM | LAN | WM | 0.286 (0.192) | 0.291 (0.190) | 0.278 (0.190) | 0.279 (0.192) | 0.293 (0.191) | 0.280 (0.191) | 0.345 (0.200) | 0.344 (0.196) | 0.337 (0.200) | 0.355 (0.204) | 0.352 (0.199) | 0.346 (0.202) |
| LAN | MOT | SOC | WM | 0.291 (0.19) | 0.309 (0.191) | 0.298 (0.192) | 0.285 (0.190) | 0.297 (0.189) | 0.282 (0.189) | 0.332 (0.194) | 0.339 (0.194) | 0.334 (0.196) | 0.329 (0.195) | 0.331 (0.190) | 0.317 (0.197) |
| REL | GAM | LAN | SOC | 0.291 (0.193) | 0.286 (0.192) | 0.283 (0.191) | 0.281 (0.192) | 0.302 (0.192) | 0.286 (0.190) | 0.354 (0.199) | 0.348 (0.196) | 0.342 (0.195) | 0.347 (0.202) | 0.356 (0.197) | 0.343 (0.198) |
| REL | LAN | SOC | WM | 0.295 (0.191) | 0.305 (0.192) | 0.286 (0.191) | 0.281 (0.191) | 0.289 (0.191) | 0.299 (0.192) | 0.347 (0.194) | 0.357 (0.200) | 0.342 (0.200) | 0.336 (0.199) | 0.343 (0.198) | 0.351 (0.198) |
| GAM | LAN | SOC | WM | 0.300 (0.190) | 0.312 (0.193) | 0.289 (0.192) | 0.280 (0.192) | 0.285 (0.191) | 0.292 (0.194) | 0.345 (0.193) | 0.352 (0.197) | 0.342 (0.199) | 0.336 (0.199) | 0.339 (0.198) | 0.35 (0.201) |

| | No GSR - ICA FIX | | No GSR - No ICA FIX | |
|---|---|---|---|---|
| Rest Conditions | LR | RL | LR | RL |
| Rest Contiguous (Day1), 11.1mins | 0.247 (0.184) | 0.270 (0.193) | 0.277 (0.179) | 0.296 (0.188) |
| Rest Concatenated - 4 scans, 11.1mins | 0.343 (0.199) | | 0.338 (0.188) | |

Table S4 continued

| GSR | | | | GSR - with task regression | | | | | | GSR - without task regression | | | | | |
|---|---|---|---|---|---|---|---|---|---|---|---|---|---|---|---|
| Task1 | Task2 | Task3 | Task4 | 1L-2R-3L-4R | 1L-2L-3R-4R | 1L-2R-3R-4L | 1R-2R-3L-4L | 1R-2L-3L-4R | 1R-2L-3R-4L | 1L-2R-3L-4R | 1L-2L-3R-4R | 1L-2R-3R-4L | 1R-2R-3L-4L | 1R-2L-3L-4R | 1R-2L-3R-4L |
| REL | GAM | MOT | WM | 0.247 (0.192) | 0.241 (0.190) | 0.240 (0.189) | 0.240 (0.189) | 0.252 (0.191) | 0.242 (0.190) | 0.332 (0.205) | 0.33 (0.206) | 0.329 (0.205) | 0.336 (0.209) | 0.345 (0.205) | 0.343 (0.206) |
| REL | GAM | SOC | WM | 0.249 (0.193) | 0.257 (0.193) | 0.247 (0.191) | 0.244 (0.191) | 0.258 (0.195) | 0.256 (0.194) | 0.351 (0.206) | 0.362 (0.207) | 0.357 (0.207) | 0.357 (0.208) | 0.369 (0.204) | 0.369 (0.206) |
| GAM | MOT | SOC | WM | 0.249 (0.192) | 0.263 (0.193) | 0.236 (0.188) | 0.236 (0.188) | 0.253 (0.193) | 0.253 (0.191) | 0.323 (0.199) | 0.335 (0.203) | 0.333 (0.203) | 0.319 (0.204) | 0.323 (0.202) | 0.329 (0.207) |
| REL | MOT | SOC | WM | 0.254 (0.192) | 0.268 (0.193) | 0.252 (0.190) | 0.251 (0.191) | 0.261 (0.193) | 0.262 (0.192) | 0.331 (0.201) | 0.339 (0.204) | 0.339 (0.204) | 0.332 (0.201) | 0.334 (0.198) | 0.333 (0.204) |
| REL | GAM | MOT | SOC | 0.261 (0.194) | 0.255 (0.192) | 0.253 (0.193) | 0.253 (0.193) | 0.264 (0.193) | 0.255 (0.193) | 0.342 (0.203) | 0.344 (0.201) | 0.333 (0.199) | 0.342 (0.205) | 0.354 (0.205) | 0.349 (0.200) |
| REL | GAM | LAN | MOT | 0.268 (0.196) | 0.264 (0.197) | 0.267 (0.198) | 0.283 (0.199) | 0.278 (0.200) | 0.270 (0.198) | 0.334 (0.201) | 0.330 (0.202) | 0.326 (0.203) | 0.352 (0.206) | 0.346 (0.204) | 0.345 (0.205) |
| GAM | LAN | MOT | WM | 0.268 (0.198) | 0.269 (0.199) | 0.258 (0.196) | 0.258 (0.196) | 0.281 (0.199) | 0.260 (0.196) | 0.318 (0.205) | 0.315 (0.205) | 0.319 (0.205) | 0.312 (0.207) | 0.327 (0.205) | 0.322 (0.206) |
| REL | LAN | MOT | WM | 0.272 (0.198) | 0.273 (0.197) | 0.259 (0.195) | 0.266 (0.197) | 0.289 (0.199) | 0.274 (0.197) | 0.322 (0.205) | 0.329 (0.205) | 0.325 (0.204) | 0.325 (0.206) | 0.339 (0.204) | 0.336 (0.206) |
| REL | GAM | LAN | WM | 0.277 (0.199) | 0.267 (0.198) | 0.277 (0.198) | 0.277 (0.198) | 0.286 (0.202) | 0.267 (0.199) | 0.357 (0.208) | 0.348 (0.209) | 0.343 (0.207) | 0.368 (0.209) | 0.368 (0.208) | 0.363 (0.207) |
| REL | LAN | SOC | WM | 0.28 (0.199) | 0.298 (0.198) | 0.276 (0.197) | 0.275 (0.199) | 0.294 (0.201) | 0.297 (0.198) | 0.341 (0.205) | 0.360 (0.206) | 0.350 (0.205) | 0.346 (0.204) | 0.358 (0.203) | 0.361 (0.204) |
| GAM | LAN | SOC | WM | 0.279 (0.201) | 0.299 (0.201) | 0.266 (0.197) | 0.266 (0.197) | 0.285 (0.200) | 0.289 (0.198) | 0.344 (0.204) | 0.356 (0.205) | 0.355 (0.206) | 0.336 (0.207) | 0.347 (0.205) | 0.358 (0.208) |
| LAN | MOT | SOC | WM | 0.281 (0.199) | 0.305 (0.199) | 0.285 (0.197) | 0.269 (0.196) | 0.282 (0.199) | 0.279 (0.196) | 0.320 (0.201) | 0.334 (0.203) | 0.332 (0.204) | 0.318 (0.202) | 0.316 (0.199) | 0.322 (0.204) |
| GAM | LAN | MOT | SOC | 0.281 (0.199) | 0.288 (0.201) | 0.274 (0.200) | 0.274 (0.200) | 0.299 (0.200) | 0.274 (0.197) | 0.338 (0.205) | 0.337 (0.202) | 0.328 (0.200) | 0.322 (0.203) | 0.344 (0.205) | 0.327 (0.202) |
| REL | LAN | MOT | SOC | 0.284 (0.198) | 0.288 (0.197) | 0.278 (0.199) | 0.278 (0.199) | 0.303 (0.198) | 0.285 (0.198) | 0.333 (0.202) | 0.340 (0.201) | 0.327 (0.199) | 0.330 (0.200) | 0.346 (0.202) | 0.337 (0.201) |
| REL | GAM | LAN | SOC | 0.292 (0.198) | 0.279 (0.199) | 0.272 (0.199) | 0.286 (0.199) | 0.301 (0.201) | 0.279 (0.201) | 0.362 (0.204) | 0.357 (0.204) | 0.340 (0.203) | 0.366 (0.204) | 0.375 (0.203) | 0.364 (0.202) |

| | GSR - ICA FIX | | GSR - No ICA FIX | |
|---|---|---|---|---|
| Rest Conditions | LR | RL | LR | RL |
| Rest Contiguous (Day1), 11.1mins | 0.283 (0.192) | 0.297 (0.203) | 0.283 (0.193) | 0.298 (0.203) |
| Rest Concatenated - 4 scans, 11.1mins | 0.360 (0.207) | | 0.335 (0.203) | |

Table S5. The averaged test-retest reliability of the edgewise connections for resting state and hybrid combinations with six different tasks preprocessed with different preprocessing options (i.e. GSR, task-regression, and ICA-FIX).

| | | | | | | With task regression | | Without task regression | |
|---|---|---|---|---|---|---|---|---|---|
| Task1 | Task2 | Task3 | Task4 | Task5 | Task6 | No GSR | GSR | No GSR | GSR |
| WM-L | MOT-L | GAM-R | REL-L | LAN-R | SOC-R | 0.299 (0.194) | 0.294 (0.203) | 0.361 (0.201) | 0.366 (0.208) |
| WM-L | MOT-L | GAM-R | REL-R | LAN-R | SOC-L | 0.302 (0.195) | 0.287 (0.204) | 0.362 (0.204) | 0.363 (0.211) |
| WM-L | MOT-L | GAM-L | REL-R | LAN-R | SOC-R | 0.302 (0.194) | 0.297 (0.204) | 0.364 (0.202) | 0.379 (0.209) |
| WM-L | MOT-R | GAM-L | REL-L | LAN-R | SOC-R | 0.306 (0.194) | 0.290 (0.203) | 0.372 (0.201) | 0.374 (0.208) |
| WM-L | MOT-R | GAM-R | REL-L | LAN-R | SOC-L | 0.307 (0.194) | 0.283 (0.201) | 0.366 (0.200) | 0.359 (0.207) |
| WM-L | MOT-R | GAM-R | REL-R | LAN-L | SOC-L | 0.307 (0.196) | 0.291 (0.202) | 0.368 (0.205) | 0.370 (0.209) |
| WM-L | MOT-R | GAM-L | REL-R | LAN-R | SOC-L | 0.308 (0.194) | 0.288 (0.203) | 0.369 (0.202) | 0.374 (0.207) |
| WM-R | MOT-L | GAM-L | REL-L | LAN-R | SOC-R | 0.306 (0.195) | 0.300 (0.204) | 0.369 (0.199) | 0.374 (0.209) |
| WM-R | MOT-L | GAM-R | REL-L | LAN-R | SOC-L | 0.307 (0.194) | 0.293 (0.205) | 0.365 (0.198) | 0.360 (0.209) |
| WM-L | MOT-L | GAM-R | REL-R | LAN-L | SOC-R | 0.31 (0.194) | 0.307 (0.203) | 0.374 (0.202) | 0.378 (0.211) |
| WM-R | MOT-L | GAM-R | REL-R | LAN-L | SOC-L | 0.310 (0.194) | 0.304 (0.205) | 0.369 (0.201) | 0.378 (0.209) |
| WM-R | MOT-L | GAM-L | REL-R | LAN-R | SOC-L | 0.310 (0.194) | 0.299 (0.205) | 0.366 (0.199) | 0.377 (0.207) |
| WM-L | MOT-R | GAM-R | REL-L | LAN-L | SOC-R | 0.312 (0.195) | 0.297 (0.202) | 0.373 (0.201) | 0.374 (0.207) |
| WM-R | MOT-L | GAM-R | REL-L | LAN-L | SOC-R | 0.312 (0.195) | 0.311 (0.204) | 0.373 (0.201) | 0.378 (0.209) |
| WM-R | MOT-R | GAM-R | REL-L | LAN-L | SOC-L | 0.312 (0.193) | 0.295 (0.202) | 0.369 (0.199) | 0.367 (0.207) |
| WM-R | MOT-R | GAM-L | REL-L | LAN-R | SOC-L | 0.315 (0.192) | 0.292 (0.203) | 0.375 (0.196) | 0.365 (0.207) |
| WM-R | MOT-R | GAM-L | REL-R | LAN-L | SOC-L | 0.317 (0.193) | 0.303 (0.205) | 0.373 (0.199) | 0.381 (0.206) |
| WM-L | MOT-R | GAM-L | REL-R | LAN-L | SOC-R | 0.319 (0.195) | 0.302 (0.205) | 0.380 (0.201) | 0.382 (0.208) |
| WM-R | MOT-L | GAM-L | REL-R | LAN-L | SOC-R | 0.318 (0.194) | 0.316 (0.205) | 0.376 (0.198) | 0.389 (0.207) |
| WM-R | MOT-R | GAM-L | REL-L | LAN-L | SOC-R | 0.320 (0.194) | 0.304 (0.205) | 0.381 (0.200) | 0.378 (0.209) |
| Rest Conditions | | | | | | No GSR - ICA FIX | GSR - ICA FIX | No GSR - No ICA FIX | GSR - No ICA FIX |
| Rest Continuous LR (Day1) | | | | | | 0.274 (0.187) | 0.315 (0.195) | 0.313 (0.182) | 0.320 (0.191) |
| Rest Continuous RL (Day1) | | | | | | 0.289 (0.193) | 0.318 (0.203) | 0.328 (0.187) | 0.319 (0.192) |
| Rest Concatenated (#1-#200 time-points from each of 4 scans, and #401-#600 from LR(Day1) and RL(Day2) scans | | | | | | 0.363 (0.203) | 0.387 (0.208) | 0.361 (0.190) | 0.37 (0.202) |
| Rest Concatenated (#1-#200 time-points from each of 4 scans, and #401-#600 from RL(Day1) and LR(Day2) scans | | | | | | 0.373 (0.203) | 0.391 (0.211) | 0.376 (0.191) | 0.375 (0.206) |

Table S6. The test-retest reliability of the edgewise connections for contiguous and concatenated resting-state data using partial correlation and full correlation based on Scheafer100 parcellation.

| Task Conditions | Partial Correlation | Full Correlation |
|---|---|---|
| | Mean ICC (std), Discriminability | Mean ICC (std), Discriminability |
| Rest Contiguous RL (Day 2)<br>15 minute segment | 0.043 (0.053), 0.63 | 0.063 (0.069), 0.73 |
| Rest Concatenated (Day 1, Day 2)<br>2 x 7.5 minute segments | 0.046 (0.055), 0.67 | 0.064 (0.069), 0.76 |

Table S7. The test-retest reliability of the edgewise connections for concatenated task conditions calculated using partial correlation and full correlation based on Scheafer100 parcellation.

| Task Conditions (2 segments, 5.6 min in total) | Partial Correlation | Full Correlation |
|---|---|---|
| | Mean ICC (std), Discriminability | |
| SOC | 0.083 (0.113), 0.720 | 0.183 (0.110), 0.924 |
| GAM | 0.082 (0.114), 0.637 | 0.130 (0.110), 0.847 |
| REL | 0.080 (0.110), 0.637 | 0.138 (0.109), 0.886 |
| MOT | 0.079 (0.111), 0.610 | 0.135 (0.111), 0.812 |
| LAN | 0.077 (0.110), 0.569 | 0.212 (0.110), 0.950 |
| Rest (Day 2) | 0.074 (0.109), 0.490 | 0.215 (0.168), 0.844 |
| WM | 0.074 (0.108), 0.512 | 0.146 (0.108), 0.878 |

Table S8. The test-retest reliability of the edgewise connections for hybrid conditions concatenated from two scans from two tasks and calculated using partial correlation and full correlation based on Scheafer100 parcellation.

| Hybrid Conditions 2 segments, 5.6 min in total | | Partial Correlation | Full Correlation |
|---|---|---|---|
| Task 1 | Task 2 | Mean ICC (std), Discriminability | |
| GAM | SOC | 0.083 (0.114), 0.739 | 0.2 (0.108), 0.906 |
| MOT | SOC | 0.083 (0.114), 0.731 | 0.19 (0.107), 0.876 |
| LAN | SOC | 0.082 (0.112), 0.672 | 0.235 (0.109), 0.942 |
| REL | SOC | 0.081 (0.114), 0.689 | 0.193 (0.109), 0.875 |
| LAN | WM | 0.081 (0.113), 0.554 | 0.218 (0.107), 0.913 |
| REL | LAN | 0.081 (0.112), 0.687 | 0.215 (0.111), 0.96 |
| GAM | LAN | 0.081 (0.112), 0.663 | 0.205 (0.111), 0.93 |
| REL | MOT | 0.08 (0.113), 0.626 | 0.168 (0.111), 0.866 |
| REST | WM | 0.079 (0.113), 0.56 | 0.189 (0.105), 0.813 |
| LAN | MOT | 0.079 (0.112), 0.605 | 0.203 (0.109), 0.922 |
| REL | GAM | 0.079 (0.111), 0.641 | 0.178 (0.111), 0.88 |
| REST | MOT | 0.079 (0.111), 0.626 | 0.181 (0.109), 0.837 |
| REST | GAM | 0.079 (0.111), 0.595 | 0.189 (0.111), 0.819 |
| GAM | MOT | 0.079 (0.111), 0.585 | 0.158 (0.111), 0.826 |
| REST | LAN | 0.078 (0.109), 0.587 | 0.223 (0.109), 0.899 |
| MOT | WM | 0.077 (0.11), 0.61 | 0.169 (0.108), 0.817 |
| SOC | WM | 0.077 (0.109), 0.572 | 0.214 (0.112), 0.924 |
| GAM | WM | 0.076 (0.109), 0.527 | 0.173 (0.109), 0.833 |
| REST | SOC | 0.076 (0.109), 0.52 | 0.217 (0.108), 0.86 |
| REL | WM | 0.076 (0.108), 0.569 | 0.192 (0.109), 0.901 |
| REST | REL | 0.076 (0.108), 0.541 | 0.196 (0.112), 0.881 |
| Rest Conditions | | | |
| Rest Contiguous (Day 2 RL) | | 0.069 (0.104), 0.492 | 0.095 (0.104), 0.994 |
| Rest Concatenated - 2 scans | | 0.078 (0.11), 0.49 | 0.212 (0.11), 0.844 |

Table S9. The averaged test-retest reliability of the edgewise connections for hybrid conditions concatenated from four scans from two tasks and calculated using partial correlation and full correlation based on Scheafer100 parcellation.

| Hybrid Conditions 4 segments, 11.1 min in total | | Partial Correlation | Pearson Correlation |
|---|---|---|---|
| Task 1 | Task 2 | Mean ICC (std), Discriminability | |
| REST1 | LAN | 0.078 (0.111), 0.958 | 0.318 (0.186), 0.995 |
| SOC | WM | 0.08 (0.112), 0.952 | 0.262 (0.177), 0.993 |
| REST1 | SOC | 0.083 (0.113), 0.95 | 0.283 (0.179), 0.994 |
| GAM | LAN | 0.082 (0.112), 0.948 | 0.297 (0.187), 0.995 |
| LAN | SOC | 0.082 (0.112), 0.947 | 0.326 (0.179), 0.995 |
| REST1 | MOT | 0.082 (0.112), 0.945 | 0.258 (0.178), 0.985 |
| LAN | MOT | 0.081 (0.112), 0.944 | 0.29 (0.184), 0.994 |
| LAN | WM | 0.078 (0.112), 0.944 | 0.299 (0.186), 0.995 |
| REST1 | GAM | 0.082 (0.114), 0.943 | 0.26 (0.181), 0.99 |
| GAM | MOT | 0.077 (0.109), 0.943 | 0.232 (0.17), 0.988 |
| REST1 | WM | 0.081 (0.113), 0.943 | 0.245 (0.179), 0.988 |
| MOT | SOC | 0.081 (0.113), 0.936 | 0.265 (0.17), 0.995 |
| REL | SOC | 0.08 (0.11), 0.936 | 0.263 (0.175), 0.994 |
| REL | LAN | 0.078 (0.108), 0.935 | 0.316 (0.188), 0.998 |
| REL | WM | 0.078 (0.109), 0.934 | 0.249 (0.182), 0.995 |
| REST1 | REL | 0.078 (0.111), 0.933 | 0.268 (0.176), 0.997 |
| GAM | WM | 0.079 (0.112), 0.932 | 0.224 (0.172), 0.991 |
| GAM | SOC | 0.078 (0.111), 0.932 | 0.268 (0.176), 0.994 |
| MOT | WM | 0.079 (0.111), 0.926 | 0.237 (0.171), 0.987 |
| REL | MOT | 0.076 (0.107), 0.919 | 0.246 (0.171), 0.993 |
| REL | GAM | 0.081 (0.112), 0.917 | 0.249 (0.176), 0.995 |
| Rest Conditions | | | |
| Rest Contiguous (Day 2 RL) | | 0.084 (0.115), 0.94 | 0.097 (0.124), 0.991 |
| Rest Concatenated - 4 scans | | 0.081 (0.114), 0.96 | 0.322 (0.181), 0.994 |

Table S10. The averaged test-retest reliability of the edgewise connections from FC calculated with partial correlation and full correlation based on Scheafer100 parcellation for Strategy-A (average multiple FC before reliability calculation) and Strategy-T (concatenate first then average) for 20 minutes and 30 minutes of total data.

| Task Conditions | Partial Correlation | | Full Correlation | |
|---|---|---|---|---|
| | Mean ICC (std), Discriminability | | | |
| | Strategy-T | Strategy-A | Strategy-T | Strategy-A |
| 20 Minutes in total (2 segments x833TR) | 0.068 (0.072), 0.841 | 0.057 (0.055), 0.643 | 0.516 (0.092), 0.971 | 0.419 (0.097), 0.954 |
| 30 Minutes in total (2 segments x1200TR) | 0.095 (0.089), 0.949 | 0.143 (0.073), 0.975 | 0.558 (0.088), 0.979 | 0.477 (0.094), 0.972 |
